# PilB from *Streptococcus sanguinis* is a bimodular type IV pilin with a direct role in adhesion

**DOI:** 10.1101/2021.01.27.428410

**Authors:** Claire Raynaud, Devon Sheppard, Jamie-Lee Berry, Ishwori Gurung, Vladimir Pelicic

## Abstract

Type IV pili (T4P) are functionally versatile filamentous nanomachines, nearly ubiquitous in prokaryotes. They are predominantly polymers of one major pilin, but also contain minor pilins whose functions are often poorly defined, and likely to be diverse. Here, we show that the minor pilin PilB from the T4P of *S. sanguinis* displays an unusual bimodular 3D structure, with a bulky von Willebrand factor A-like (vWA) module “grafted” onto a small pilin module via a short unstructured loop. Structural modelling suggests that PilB is only compatible with a localisation at the tip of T4P. By performing a detailed functional analysis, we found that (i) the vWA module contains a canonical metal ion-dependent adhesion site (MIDAS), preferentially binding Mg^2+^ and Mn^2+^, (ii) abolishing metal-binding has no impact on the structure of PilB or piliation, (iii) metal-binding is important for *S. sanguinis* T4P-mediated twitching motility and adhesion to eukaryotic cells, and (iv) the vWA module shows an intrinsic binding ability to several host proteins. These findings reveal an elegant, yet simple, evolutionary tinkering strategy to increase T4P functional versatility, by grafting an adhesive module onto a pilin for presentation by the filaments. This strategy appears to have been extensively used by bacteria, in which modular pilins are widespread and exhibit an astonishing variety of architectures.

## INTRODUCTION

Type IV pili (T4P) are functionally versatile filaments widespread in prokaryotes, implicated in a variety of functions such as adhesion, twitching motility, DNA uptake *etc*^1^. T4P are helical polymers consisting of type IV pilins, usually one major pilin and several minor (low abundance) ones, assembled by distinctive multi-protein machineries. These defining features are shared by a superfamily of filamentous nanomachines known as type IV filaments (T4F)^1^, which are ubiquitous in prokaryotes^2^.

T4P have been intensively studied for decades in diderm bacteria because they play a central role in pathogenesis in several important human pathogens^3^. The following global picture of T4P biology has emerged from these studies. The pilus subunits, type IV pilins, are characterised by a short N-terminal sequence motif known as class III signal peptide, which consists of a hydrophilic leader peptide ending with a small residue (Gly or Ala), followed by a tract of 21 predominantly hydrophobic residues^4^. This tract constitutes the N-terminal segment (α1N) of an α-helix (α1) of ∼50 residues, which is the universally conserved structural feature in type IV pilins. Usually, the α1N helix protrudes from a globular head most often consisting of a β-sheet composed of several antiparallel β-strands, which gives pilins their characteristic “lollipop” shape^4^. The hydrophilic leader peptide is then processed by a dedicated prepilin peptidase^5^ upon pilin translocation across the cytoplasmic membrane (CM) by the general secretory pathway^6,7^. Processed pilins remain embedded in the CM via their α1N, generating a pool of subunits ready for polymerisation. Filament assembly, which occurs from tip to base, is mediated at the CM by a complex multi-protein machinery (10-20 components)^1^, centred on an integral membrane platform protein and a cytoplasmic extension ATPase^8^. Recent cryo-EM structures have revealed that T4P are right-handed helical polymers in which pilins are held together by extensive interactions between their α1N helices, which are partially melted and run approximately parallel to each other within the filament core^9,10^. One of the properties of T4P key for their functional versatility is their ability to retract, which has been best characterised for T4aP (where “a” denotes the subtype). In T4aP, retraction results from rapid filament depolymerisation powered by the cytoplasmic retraction ATPase PilT^11^, which generates important tensile forces^12,13^.

Studying T4P in monoderm bacteria represents a promising alternative research avenue^14^. *Streptococcus sanguinis*, a commensal of the oral cavity that commonly causes life-threatening infective endocarditis (IE), has emerged as a monoderm model for deciphering T4P biology^15^. Our comprehensive functional analysis of *S. sanguinis* T4P^16^ revealed that they are canonical T4aP. Indeed, filaments are (i) assembled by a multi-protein machinery similar to diderm T4aP species, but simpler with only ten components, (ii) retracted by a PilT ATPase, generating tensile forces similar to diderm species, and (iii) powering intense twitching motility, leading to spreading zones around bacteria growing on plates, visible by the naked eye. Subsequently, we performed a global biochemical and structural analysis of *S. sanguinis* T4P^17^, showing that (i) they are hetero-polymers composed of two major pilins, PilE1 and PilE2, rather than one as normally seen, (ii) the major pilins display classical type IV pilin 3D structure, and (iii) the filaments contain a low abundance of three minor pilins (PilA, PilB, and PilC), which are required for piliation.

The present study was prompted by a perplexing observation, *i*.*e*., that the minor pilin PilB harbours a protein domain that has been extensively studied in several eukaryotic proteins where it mediates adhesion to a variety of protein ligands^18^. This suggested that PilB might be an adhesin, promoting T4P-mediated adhesion of *S. sanguinis* to host cells and proteins. Therefore, since both the molecular mechanisms of T4P-mediated adhesion and the exact role of minor pilins in T4P biology remain incompletely understood^1^, we decided to perform a structure/function analysis of PilB, which is reported here. This uncovered a widespread strategy for minor pilins to enhance the functional properties of T4P.

## RESULTS

### PilB displays a modular pilin architecture

PilB, one of the three minor pilins in *S. sanguinis* T4P^17^, exhibits a canonical N-terminal class III signal peptide, the defining feature of type IV pilins^4^. This sequence motif consists of a seven-residue leader peptide composed predominantly of hydrophilic and neutral amino acids (aa), ending with a conserved Gly (Fig. 1A). This leader peptide, which is processed by the prepilin peptidase PilD^17^, is followed by a stretch of 21 predominantly hydrophobic aa, except for a negatively charged Glu in position 5 (Fig. 1A). Processed PilB is unusually large for a pilin, with a predicted molecular mass of 50.5 kDa (Fig. 1B). For comparison, the two major pilins of *S. sanguinis* T4P, PilE1 and PilE2^16^, have typical pilin sizes of 14.7 and 14.1 kDa, respectively (Fig. 1B). The larger size of PilB is due to the presence of a C-terminal domain (Fig. 1B) readily detectable by a bioinformatic analysis, which belongs to the von Willebrand factor A-like domain superfamily (InterPro entry IPR036465). We will refer to this domain as vWA. The prototypical vWA domain is found in the von Willebrand factor (vWF), a human blood protein required for haemostasis^19^, the physiological process that prevents/stops bleeding. vWA domains, which are found in more than 300,000 proteins in the three domains of life, have been extensively studied in eukaryotic proteins where they mediate adhesion to a variety of protein ligands^18^. They have been much less studied in bacteria. Of note, the PilB vWA domain is predicted to contain a metal coordination site known as MIDAS, for metal ion-dependent adhesion site^20^ (Fig. 1A), which was found to be important for ligand-binding in several eukaryotic vWA-containing proteins^21^.

**Fig. 1.**
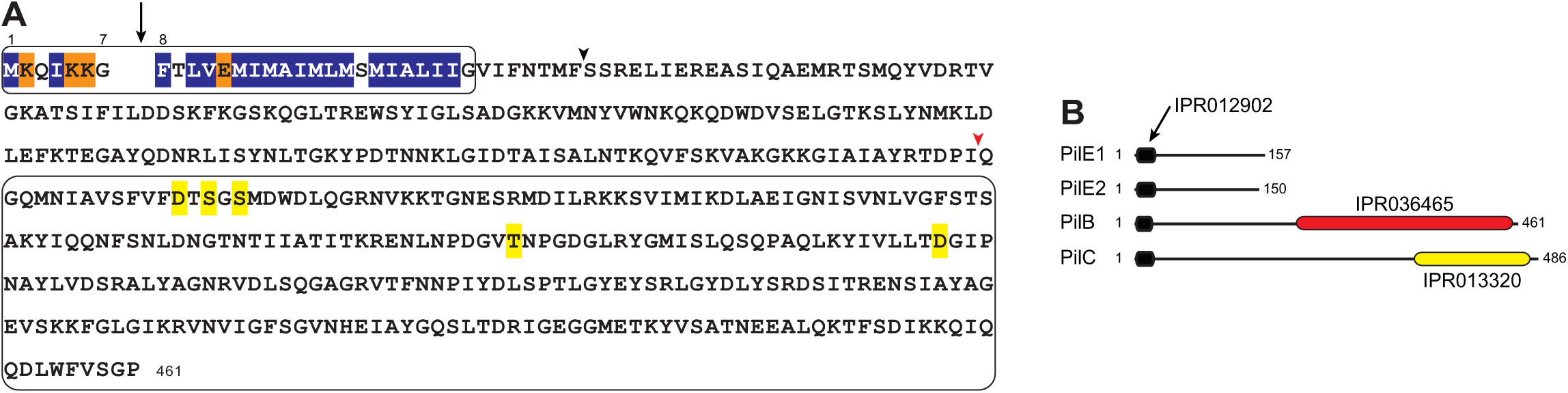
Bioinformatic analysis of PilB. **A**) Relevant features of PilB. The sequence is from *S. sanguinis* 2908. The N-terminal class III signal peptide, the defining feature of type IV pilins, is boxed. The 7-aa long leader peptide contains mostly hydrophilic (shaded in orange) and neutral (no shading) residues, and it ends with a conserved Gly. This leader peptide is processed by the prepilin peptidase PilD, which is indicated by the vertical arrow, generating a protein of 454 residues (50.5 kDa). The processed protein starts with a tract of 21 predominantly hydrophobic residues (shaded in blue), which invariably form an extended α-helix that is the main assembly interface within filaments. The C-terminal vWA module (IPR002035) in PilB is boxed, with the conserved residues forming the MIDAS highlighted in yellow. Arrowheads indicate the proteins that were produced and purified in this study, consisting of either two modules (black arrowhead) or just the vWA module (red arrowhead). **B**) Modular architectures of PilB and PilC minor pilins compared to the major pilins PilE1/PilE2. The proteins, from *S. sanguinis* 2908, have been drawn to scale. The black rounded rectangles correspond to the IPR012902 motif that is part of the class III signal peptide. The C-terminal domains in PilB and PilC are highlighted by coloured rounded rectangles, vWA domain in PilB (red) and lectin domain in PilC (yellow).

The above-described architecture is unusual for type IV pilins for two reasons. First, in contrast to classical pilins that consist only of a pilin module^4^, defined by a short N-terminal IPR012902 motif within the class III signal peptide (Fig. 1B), PilB apparently has an additional module. Second, the extra C-terminal module in PilB corresponds to a well-defined functional domain not specific to T4P biology, vWA, which is often associated with adhesion to protein ligands^18,21^. This has not been previously reported in T4P. This is what we call a modular architecture, and why we refer to PilB as a modular pilin.

Taken together, these findings suggest that PilB is a modular pilin in which a functional module has been grafted during evolution onto a pilin moiety in order to promote T4P-mediated adhesion of *S. sanguinis* to protein ligands.

### Crystal structure of PilB reveals a bimodular pilin in which a small type IV pilin module is linked to a bulky vWA module via a short loop

High-resolution structural information is required to confirm that PilB is composed of two modules, but also to understand how modular pilins are polymerised in the filaments and how they modulate T4P functionality. We therefore endeavoured to solve the 3D structure of PilB by X-ray crystallography. To facilitate protein purification, we used a synthetic *pilB* gene codon-optimised for expression in *Escherichia coli*, and produced a recombinant protein in which the N-terminal 35 aa of PilB (encompassing the hydrophobic α1N) (Fig. 1A) were replaced by a hexahistidine tag (6His)^17^. This is a commonly used approach in the field since the truncation of α1N has minimal structural impact on the rest of the protein^22^. The resulting 48.4 kDa 6His-PilB protein was soluble and could be purified using a combination of affinity and gel-filtration chromatography. The protein readily crystallised in multiple conditions, and after optimising the best diffracting crystals, we collected a complete dataset on crystals forming in the space group *P*6_1_ (Table 1). After phase determination, done using crystals produced in the presence of seleno-methionine (SeMet), we solved the 2.26 Å structure of native 6His-PilB. As can be seen in Fig. 2A, this structure reveals a clear bimodular architecture with a small pilin moiety (highlighted in blue) linked to a bulky vWA moiety (in red) by a short nine-residue unstructured loop (grey).

**Table 1.**
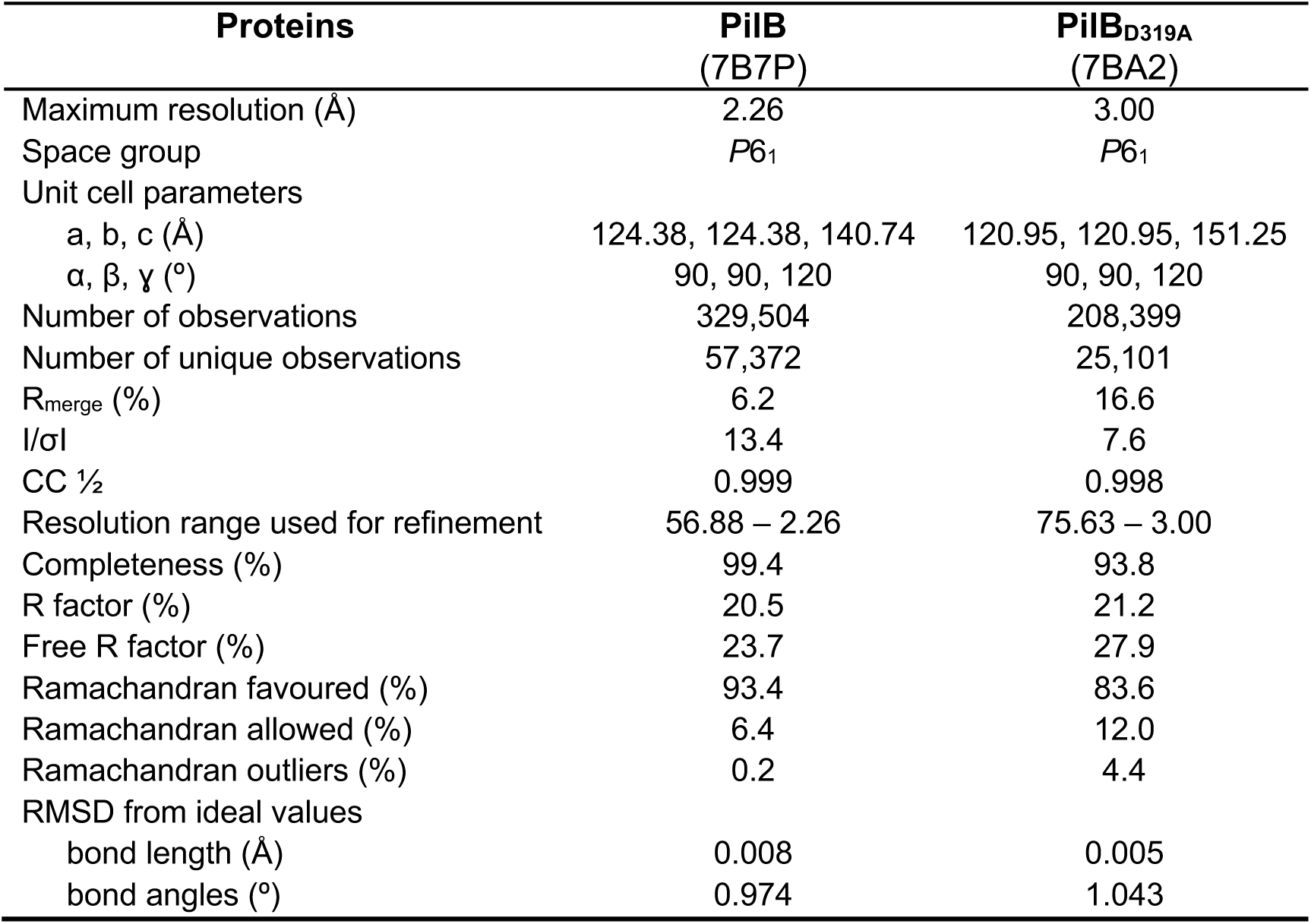
Crystal structures data collection and refinement statistics.

| **Proteins** | **PilB**<br>**(7B7P)** | **PilB<sub>D319A</sub>**<br>**(7BA2)** |
| --- | --- | --- |
| Maximum resolution (Å) | 2.26 | 3.00 |
| Space group | *P*6<sub>1</sub> | *P*6<sub>1</sub> |
| Unit cell parameters |  |  |
| a, b, c (Å) | 124.38, 124.38, 140.74 | 120.95, 120.95, 151.25 |
| α, β, γ (°) | 90, 90, 120 | 90, 90, 120 |
| Number of observations | 329,504 | 208,399 |
| Number of unique observations | 57,372 | 25,101 |
| R<sub>merge</sub> (%) | 6.2 | 16.6 |
| I/σI | 13.4 | 7.6 |
| CC ½ | 0.999 | 0.998 |
| Resolution range used for refinement | 56.88 – 2.26 | 75.63 – 3.00 |
| Completeness (%) | 99.4 | 93.8 |
| R factor (%) | 20.5 | 21.2 |
| Free R factor (%) | 23.7 | 27.9 |
| Ramachandran favoured (%) | 93.4 | 83.6 |
| Ramachandran allowed (%) | 6.4 | 12.0 |
| Ramachandran outliers (%) | 0.2 | 4.4 |
| RMSD from ideal values |  |  |
| bond length (Å) | 0.008 | 0.005 |
| bond angles (°) | 0.974 | 1.043 |

**Fig. 2.**
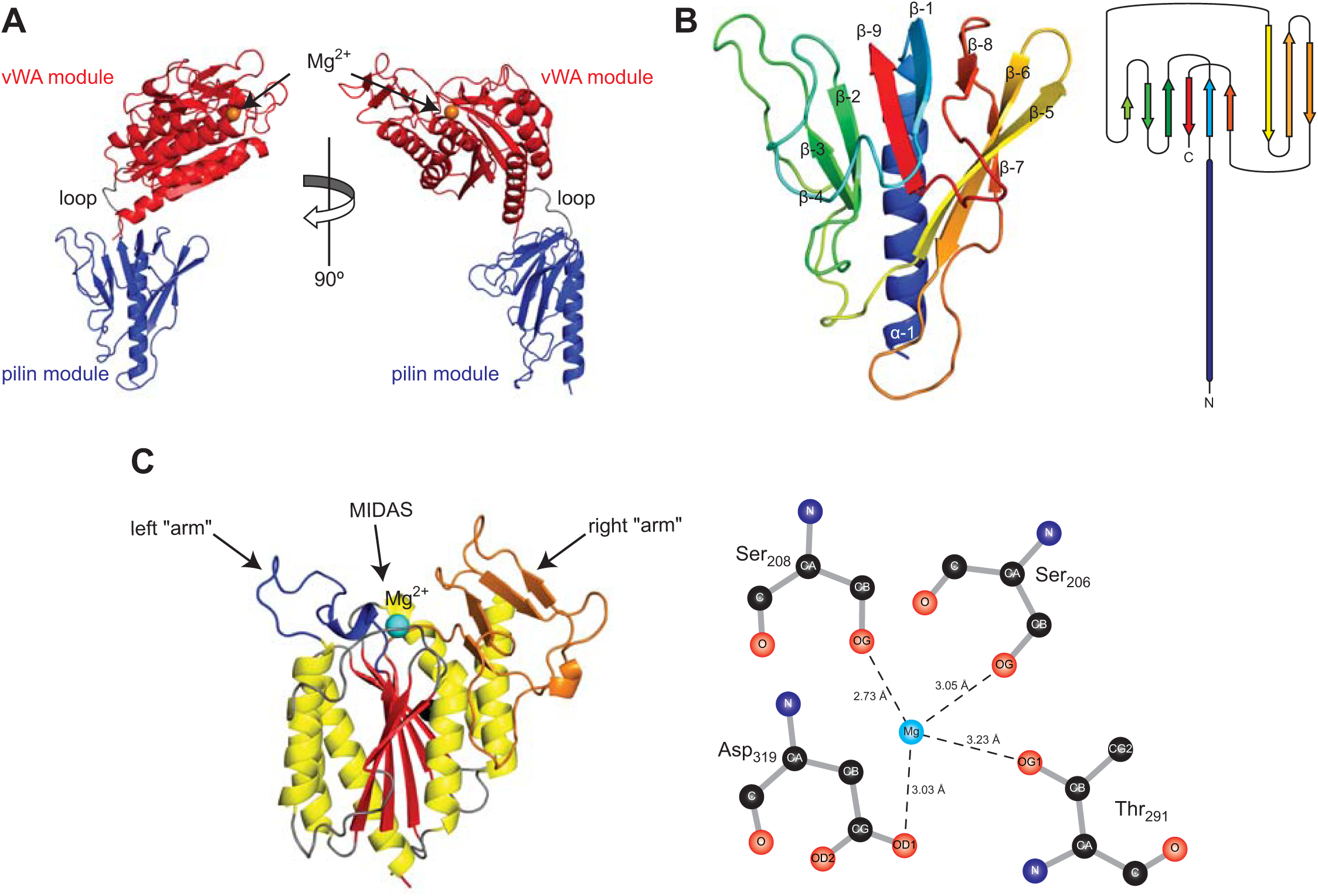
Crystal structure of PilB. **A**) Orthogonal cartoon views of the 6His-PilB structure in which the two distinct modules have been highlighted in blue (pilin module) and red (vWA module), while the short loop connecting them is in grey. The orange sphere represents a magnesium ion. **B**) Left, close-up cartoon view of the pilin module coloured in rainbow spectrum from blue (N-terminus) to red (C-terminus). Right, topology diagram of the pilin module structure. **C**) Left, close-up cartoon view of the vWA module in which the β-strands composing the central β-sheet are highlighted in red, while the surrounding α-helices are highlighted in yellow. The connecting loops are in grey, except for the two “arms” on top of the structure (coloured in orange and blue), which surround the MIDAS. Right, diagram of the magnesium coordination by the conserved MIDAS residues in the vWA module of PilB. Coordinating oxygen atoms are shown with dashed lines corresponding to hydrogen bonds.

While a bioinformatic analysis could only predict that the extreme N-terminus of PilB corresponds to an IPR012902 class III signal peptide motif, our structure reveals that the first 180 residues of processed PilB clearly display a type IV pilin fold^4^ and thus indeed correspond to a pilin module (Fig. 2B). The pilin module exhibits a long N-terminal α-helix packed against, not one β-sheet as usual, but two consecutive β-sheets consisting of six and three β-strands respectively, which together form the globular head of the pilin. The 432918 topology, *i*.*e*., the order of the β-strands in the first β-sheet (Fig. 2B), is unusual since the β-strands are not contiguous along the protein sequence. Moreover, the last portion of this β-sheet forms a Ψ-loop^23^ in which two antiparallel strands (β8 and β9) are linked via β1 in between, connected to both of them by hydrogen bonds. This motif occurs rarely in proteins^23^.

As for the vWA module (Fig. 2C), the structure strengthens the predictions of the bioinformatic analysis. The vWA moiety of PilB adopts a canonical vWA fold^20,24^, with a central β-sheet (composed of five parallel and one antiparallel β-strands) surrounded on both sides by a series of α-helices. Consequently, the vWA module of PilB shows high structural similarity to many vWA-containing proteins with which it shares little sequence identity. For example, the vWA module of PilB is very similar to the third vWA domain of human vWF^25^ (Fig. S1), with a root mean square deviation (RMSD) of 1.72 Å when the two structures are superposed. As in eukaryotic vWA-containing proteins^20,24^, PilB exhibits a MIDAS located on top of the central β-sheet (Fig. 2C). However, in contrast to these proteins, the MIDAS in PilB is flanked by two protruding “arms”, which is reminiscent of the RrgA adhesin from *Streptococcus pneumoniae*^26^. The first arm is mainly unstructured, while the second folds into a four-stranded β-sheet (Fig. 2C). The MIDAS motif in PilB, which is formed by residues conserved in vWA-containing proteins, non-contiguous in the sequence (Fig. 1A) but in close proximity in the 3D structure (Fig. 2C), is functional since it coordinates a metal ion in the crystal. We have modelled the metal as Mg^+2^ because of its abundance in the protein expression medium and the high affinity of PilB for it (see below). The Ser_206_, Ser_208_, Thr_291_ and Asp_319_ residues in the MIDAS motif^20^ of PilB form direct hydrogen bonds with the metal through oxygen atoms (Fig. 2C), while two additional coordination sites are provided by water molecules.

An important biological implication of the PilB structure is that modular pilins, despite their large size, are likely to be polymerised into T4P in the same way as classical pilins^4^, *i*.*e*., via their N-terminal pilin module. We therefore tested by structural modelling whether PilB could pack into filaments. First, we produced a full-length 3D structural model of PilB including the missing α1N (Fig. S2), which was truncated in the recombinant protein that we purified. Since a portion of α1N in major pilins is melted during filament assembly, as observed in several T4aP cryo-EM structures^9,10^, the α1N of PilB was modelled with a melted segment. This is consistent with the presence of the helix-breaking Gly residue in position 21 of α1N (Fig. 1A). Then, we fitted this full-length PilB into a previously generated model of *S. sanguinis* T4P, a right-handed helical heteropolymer where major pilins PilE1/PilE2 are held together by interactions between their α1N helices (Fig. 3A), which was based on the cryo-EM structure of *Neisseria meningitidis* T4P^9^. Despite its unusual modular structure, PilB can be readily modelled into T4P, its pilin module establishing extensive hydrophobic interactions via its α1N with the α1N of neighbouring major pilins (Fig. 3A). This suggests that PilB will assemble into filaments in the same way as classical pilins^9,10^. However, PilB can only be accommodated at the tip of the filaments because the bulky vWA module sits on top of the pilin module in the PilB structure, and essentially prevents other pilin subunits from being modelled above it (Fig. 3B). Accordingly, when PilB is modelled in the body of the filament (Fig. S3A), it exhibits important steric clashes with neighbouring major pilins (Fig. S3B).

**Fig. 3.**
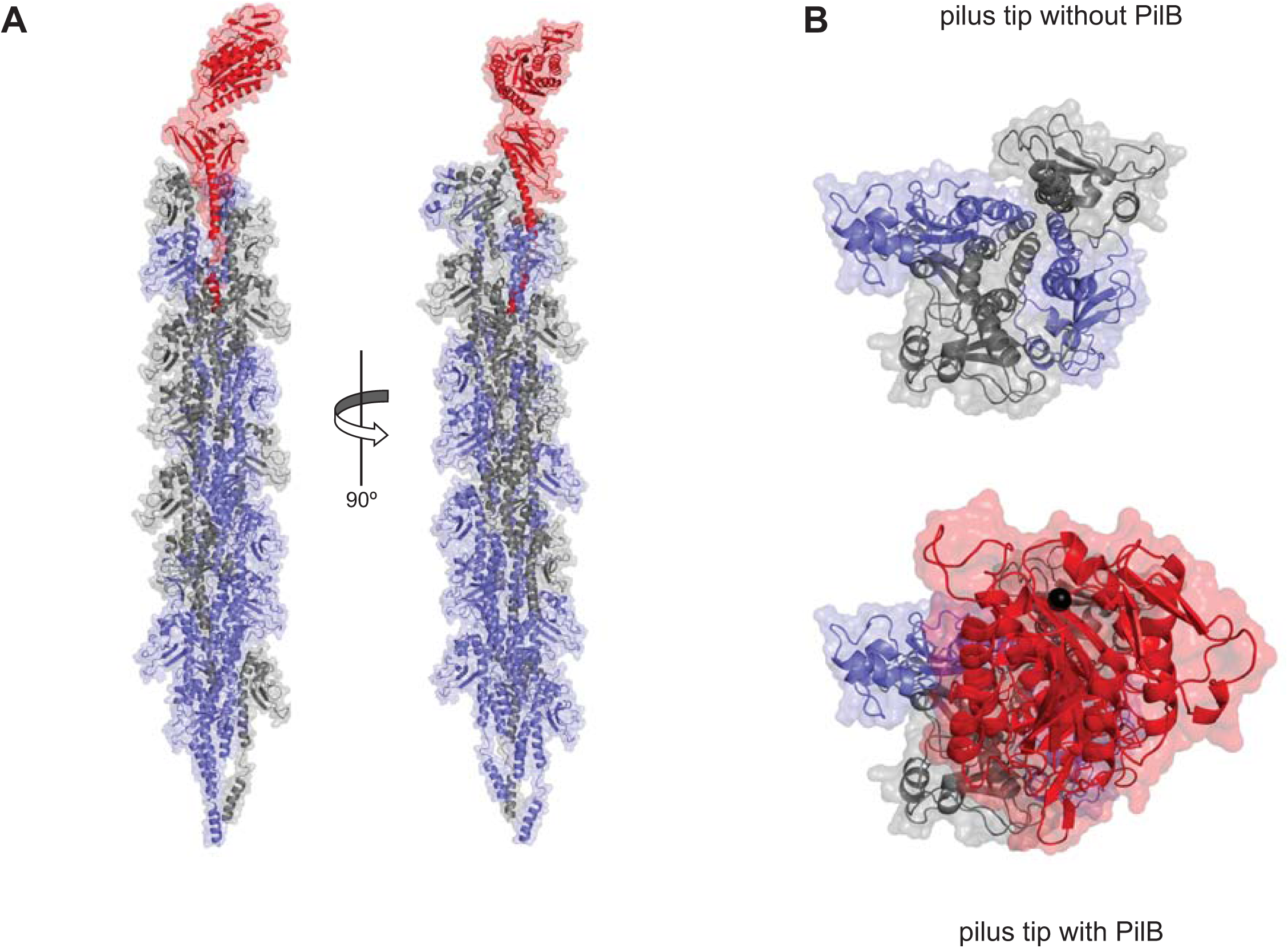
3D model of PilB in *S. sanguinis* T4P. **A**) Packing of PilB (red) into *S. sanguinis* T4P, which is a right-handed helical heteropolymer of two major pilins PilE1 (blue) and PilE2 (grey). **B**) View of the T4P tip capped by PilB, or not.

Together, these structural findings show that PilB is a bimodular protein composed of two fused but clearly distinct structural modules. The pilin module adopts a canonical type IV pilin fold^4^, which explains how modular pilins are polymerised into T4P, most probably at their tip. The second module, which is linked to the end of the pilin module via a short unstructured loop, adopts a vWA fold^20,24^ with a clearly defined MIDAS that coordinates a metal. Since the vWA motif in many eukaryotic proteins is involved in adhesion to protein ligands^18,21^, our structure strengthens our working hypothesis that PilB might be an adhesin.

### Functional analysis of the MIDAS in PilB reveals that metal binding, although structurally dispensable, is important for T4P functionality

Our PilB structure revealed that Mg^2+^, despite not being added during crystallisation, is bound by the MIDAS. In eukaryotic proteins, the MIDAS sometimes coordinates Mn^2+^ as well^20,24^. We therefore tested the metal binding specificity of the MIDAS in PilB using ThermoFluor. This fluorescent-based method, which measures changes in thermal denaturation temperature, is a commonly used approach for detecting and quantifying protein-ligand interactions^27^. We determined the affinity of purified PilB for the Mg^2+^, Mn^2+^, and Ca^2+^ divalent cations (Fig. 4A). While no binding was detected to Ca^2+^, we found that PilB binds Mg^2+^ and Mn^2+^ efficiently in the micromolar range, with estimated Kd of 70 and 54 µM, respectively. To confirm that metal binding involves the MIDAS motif, we produced the PilB_D319A_ protein in which the key MIDAS residue Asp_319_ (Fig. 2C) was changed into an Ala by site-directed mutagenesis. Binding assays performed with PilB_D319A_ showed that changing this one residue abolishes the metal-binding ability of PilB for both Mg^2+^ and Mn^2+^ (Fig. 4B). These findings show that the MIDAS in PilB is functional and preferentially binds Mg^2+^ and Mn^2+^.

**Fig. 4.**
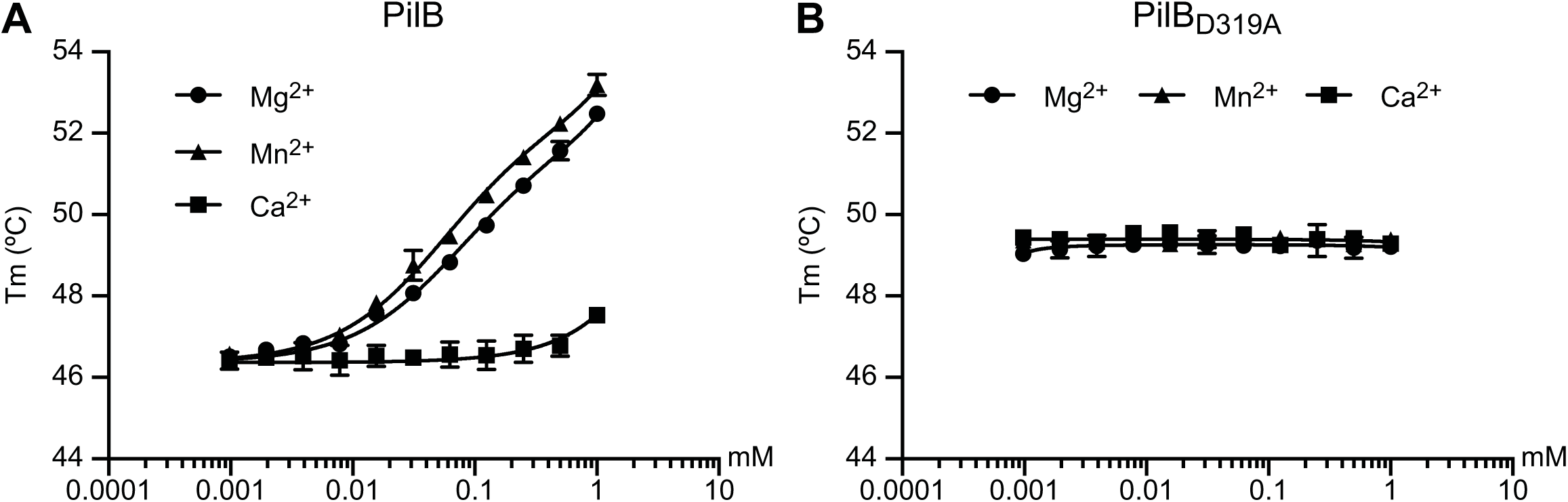
Metal binding by PilB. Purified PilB was incubated with increasing concentrations of divalent ions (Ca^2+^, Mg^2+^, Mn^2+^) and binding was quantified by ThermoFluor. **A**) Metal binding by PilB. **B**) Metal binding by PilB_D319A_, with an inactive MIDAS module. Results are the average ± standard deviations from 3 independent experiments.

Next, to determine whether metal presence/absence might impact the 3D structure of PilB, we solved the structure of PilB_D319A_ by X-ray crystallography. The PilB_D319A_ protein readily crystallised in the same condition as the wild-type (WT) protein, and we collected a complete dataset on crystals diffracting to a resolution of 3 Å (Table 1). The structure of PilB_D319A_ (Fig. 5A), which was solved by molecular replacement, shows that, no metal is occupying the mutated MIDAS pocket on top of the central β-sheet (Fig. 5B), which is consistent with results of metal binding assays. When the structures of PilB and PilB_D319A_ were compared, we found that they are essentially identical, superposing onto each other (Fig. 5C) with an RMSD of merely 0.48 Å, including the two arms flanking the MIDAS pocket. This shows that metal-binding by the MIDAS has no detectable structural impact on PilB.

**Fig. 5.**
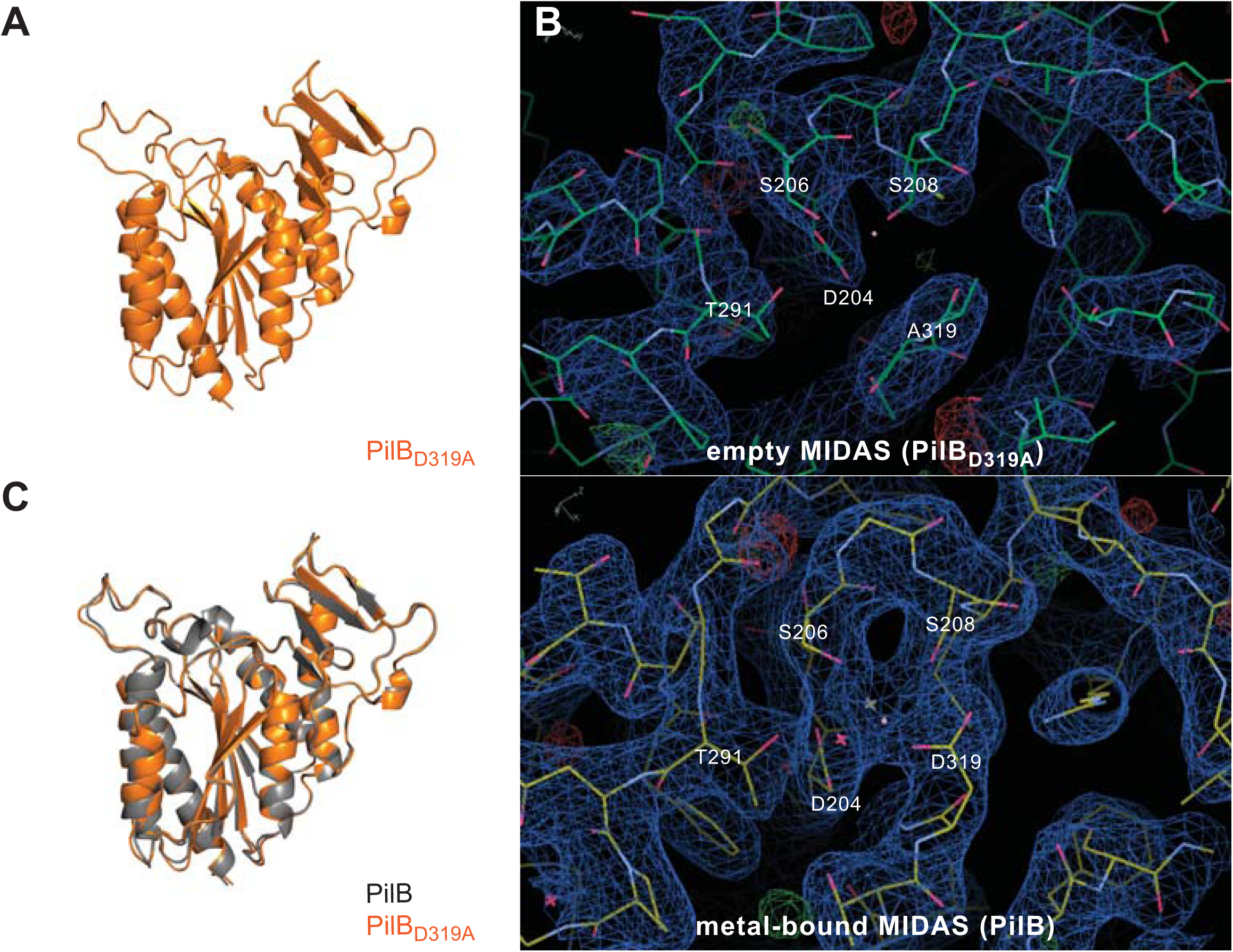
3D crystal structure of PilB_D319A_. **A**) Close-up cartoon view of the vWA module in PilB_D319A_. **B**) Comparison of electron density maps in the MIDAS pocket for the PilB_D319A_ (upper panel) and PilB (lower panel) structures. **C**) Superposition of the vWA modules of PilB (grey) and PilB_D319A_ (orange). The two structures superpose with an RMSD of 0.48 Å.

Next, we explored whether MIDAS-mediated metal-binding by PilB is important for piliation and/or T4P-powered twitching motility, both of which were previously shown to be abolished in a *ΔpilB* mutant^16^. We therefore constructed an unmarked *S. sanguinis* mutant in which the endogenous *pilB* gene was altered by site-directed mutagenesis to produce PilB_D319A_ with an inactive MIDAS. We first tested whether the *pilB*_*D319A*_ mutant retains the ability to assemble T4P using filament purification^16^. As can be seen in Fig. 6A, in which purified T4P were separated by SDS-PAGE and stained with Coomassie blue, the *pilB*_*D319A*_ mutant is piliated. This is evidenced by the presence of the two bands corresponding to major pilins PilE1 and PilE2, which are absent in a non-piliated *ΔpilD* control (Fig. 6A). Moreover, the amount of T4P that can be purified from the *pilB*_*D319A*_ mutant and WT strain appear comparable. We then tested whether the pili in the *pilB*_*D319A*_ mutant are able to mediate twitching motility^16^. For the WT strain, twitching motility is evidenced by spreading zones around bacteria grown on agar (Fig. 6B). Spreading zones were absent for the *pilB*_*D319A*_ mutant, which therefore exhibits no detectable twitching motility (Fig. 6B). This shows that the MIDAS-mediated metal-binding ability of PilB, while dispensable for piliation, is important for T4P-mediated twitching motility.

**Fig. 6.**
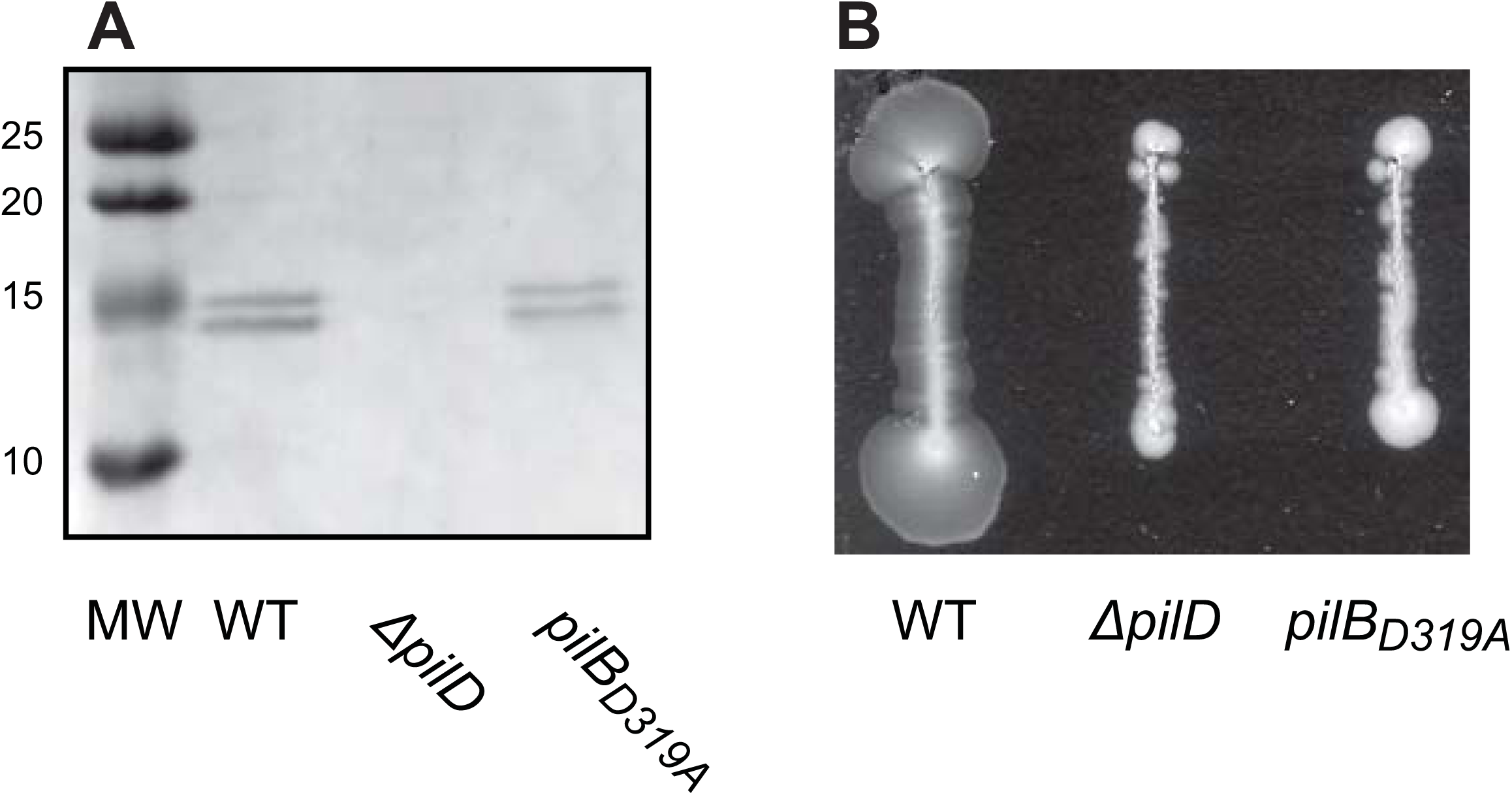
Phenotypic characterisation of a *S. sanguinis* mutant expressing PilB_D319A_ with an inactive MIDAS. **A**) Piliation was quantified by purifying T4P from cultures adjusted to the same OD_600_, using a shearing/ultracentrifugation procedure. Purified T4P (identical volumes were loaded in each lane) were separated by SDS-PAGE and stained with Coomassie blue. A molecular weight marker (MW) was run in the first lane. Molecular weights are indicated in kDa. **B**) Twitching motility was assessed by a macroscopic motility assay. Bacteria were streaked on plates, which were incubated several days at 37°C in a humid atmosphere and then photographed. Twitching motility is characterised by spreading zones around colonies.

Together, these findings show that the MIDAS in PilB is a functional metal-binding site, dispensable for piliation and protein folding, but essential for T4P functionality.

### T4P-mediated adhesion to eukaryotic cells requires PilB, which specifically binds several human proteins

Since vWA is involved in adhesion in many eukaryotic proteins^18,21^, our original hypothesis was that PilB might mediate *S. sanguinis* adhesion to host cells and/or proteins, which we aimed to test next. First, we determined whether *S. sanguinis* T4P might be involved in its well-known ability to adhere to host cells^28^. After testing a few eukaryotic cell lines, we opted for CHO cells because the WT strain adheres very efficiently to them. When CHO cells were infected by *S. sanguinis* at a multiplicity of infection (MOI) of 10, 31.6 ± 9.1 % of the bacterial inoculum adhered to the cells. In contrast, a non-piliated *ΔpilD* mutant showed a significantly reduced adhesion, with an 18-fold decrease relative to the WT (Fig. 7). Next, we tested our original assumption that PilB might be an adhesin, by quantifying the adhesion of the *pilB*_*D319A*_ mutant. As can be seen in Fig. 7, although the *pilB*_*D319A*_ mutant is piliated, its adhesion to CHO cells is dramatically impaired, with a 33-fold decrease when compared to the WT. These findings show that *S. sanguinis* T4P are multi-functional filaments also important for adhesion to eukaryotic cells, and that PilB plays an important role.

**Fig. 7.**
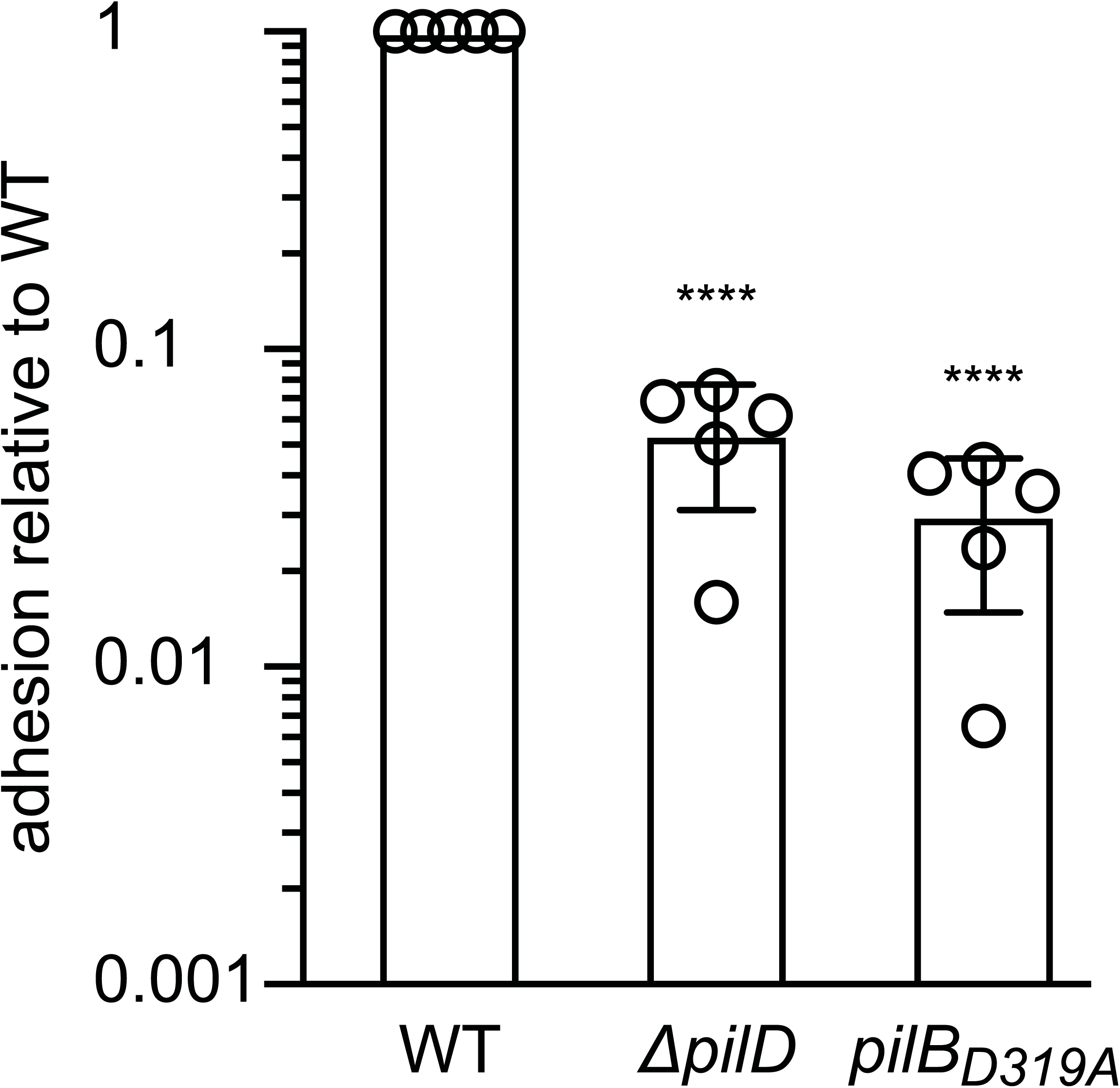
Adhesion of *S. sanguinis* to eukaryotic cells : testing the importance of T4P and the role of the MIDAS motif in PilB. Bacteria were incubated with CHO cells (MOI 10) for 1 h. After removing non-adherent bacteria by several washes, bacteria adhering to cells were enumerated by performing CFU counts. Results are expressed as adhesion relative to WT (set to 1), and are the average ± standard deviations from five independent experiments. For statistical analysis, one-way ANOVA followed by Dunnett’s multiple comparison tests were performed (\*\*\*\**P* < 0.0001).

Since the vWA domain in multiple eukaryotic proteins has been shown to mediate cell-extracellular matrix (ECM) interactions^21^, we reasoned that PilB might recognise similar ligands because it exhibits a canonical vWA module (Fig. S1). We tested this hypothesis by performing binding assays with purified PilB using enzyme-linked immunosorbent assay (ELISA). In brief, we coated 96-well plates with selected putative ligands, added serial dilutions of purified 6His-PilB, and detected binding using an anti-6His antibody. We tested binding to fibrinogen and the ECM proteins fibronectin, elastin, and laminin. While PilB exhibits no binding to BSA that was used as a negative control (Fig. 8A), dose-dependent binding to fibronectin and fibrinogen was observed, but not to the other ECM proteins that were tested (elastin and laminin). Specific binding to fibronectin and fibrinogen was in the high nanomolar range, with calculated Kd of 865 and 494 nM, respectively (Fig. 8A). Under these *in vitro* experimental conditions metal-coordination by the MIDAS is dispensable for binding to fibronectin and fibrinogen, as demonstrated by binding assays using purified PilB_D319A_ protein, which showed that PilB_D319A_ binds these ligands as well as PilB (Fig. S4). Finally, to confirm the prediction that binding of PilB to the above ligands is mediated by its vWA module, we then produced and purified the PilB_vWA_ protein corresponding only to the vWA module (see Fig. 1A). We found that PilB_vWA_ could also bind to fibronectin and fibrinogen (Fig. 8B), with calculated Kd of 337 and 997 nM, respectively, which were comparable to PilB. These findings confirm that the adhesive ability of PilB is due to its vWA module.

**Fig. 8.**
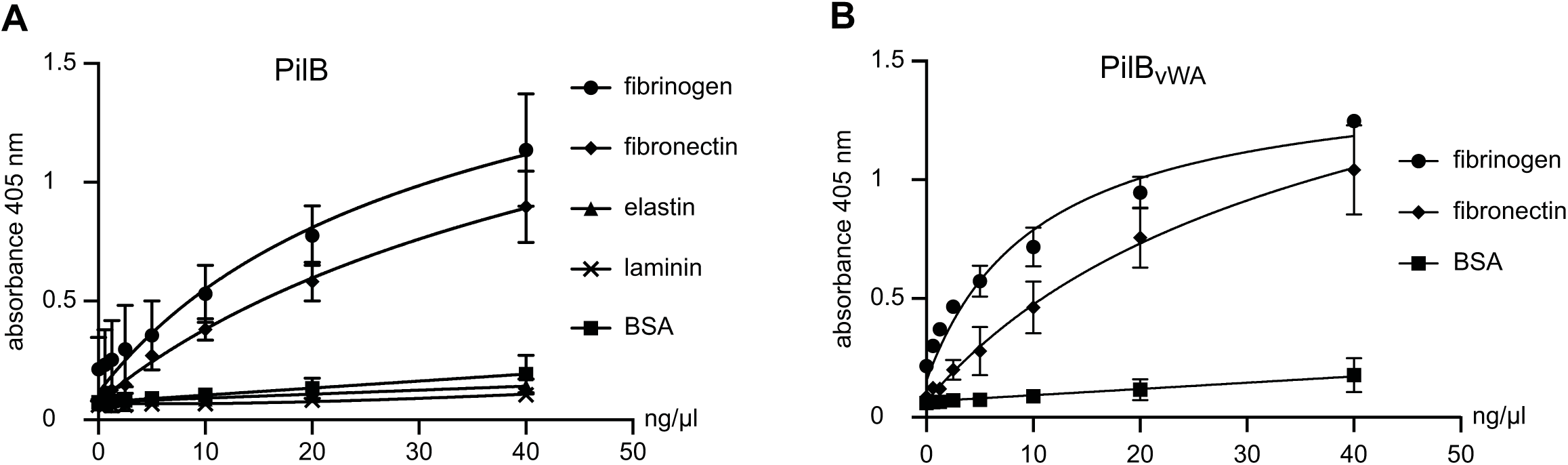
Dose-dependent binding of PilB to various protein ligands. Increasing concentrations of purified PilB was added to constant concentrations of immobilised putative ligands, and binding was quantified by ELISA. BSA served as negative control. Results are the average ± standard deviations from at least three independent experiments. **A**) Binding of PilB to fibrinogen, fibronectin, elastin, and laminin. **B**) Binding of PilB_vWA_, consisting only of the vWA module, to fibrinogen and fibronectin.

Taken together, these findings show that *S. sanguinis* T4P are multi-functional filaments mediating adhesion to eukaryotic cells, and that PilB is a *bona fide* adhesin using its vWA module to bind several human protein ligands, which it shares with eukaryotic vWA-containing proteins.

### Pilins with modular architectures are widespread in bacteria

While PilB orthologs are ubiquitous in *S. sanguinis*, which also produces a second modular pilin, PilC^17^, where the extra module belongs to the concanavalin A-like lectin/glucanase domain superfamily (IPR013320) (Fig. 1B), we wondered how widespread and how diverse modular pilins might be. We therefore searched the InterPro database^29^ for all the proteins with an N-terminal IPR012902 domain, which also contain an extra domain not specific to T4P biology. This showed that modular pilins are (i) widespread with more than 1,200 proteins displaying such architecture (Supplementary data 1), (ii) present both in monoderm and diderm species, and (iii) diverse, with as many as 264 different architectures detected. Although a bimodular architecture is the most prevalent, there are modular pilins with multiple additional domains, the most extreme case being an 860-residue protein from *Candidatus* Falkowbacteria, with 12 copies of the IPR013211 motif of unknown function (Supplementary data 1). A closer inspection of the 15 most frequent modular pilin architectures offers a glimpse of their diversity (Fig. 9). While in many of these proteins the extra domain has no clear function (IPR007001, IPR011871, IPR026906, PF05345, IPR006860, IPR003961, IPR021556), for others a function can be predicted. These functions include (i) binding to carbohydrates via PF13385 (that overlaps with the IPR013320 lectin domain superfamily), PF13620 (carbohydrate-binding-like fold), or IPR011658 (PA14 carbohydrate-binding domain), (ii) peptidase activity via IPR030392, and (iii) binding to proteins via IPR002035 (that overlaps with the IPR036465 vWA domain superfamily). These findings suggest that the rather simple modular design strategy – during which a functional module has been grafted during evolution onto a pilin moiety – appears to have been used often during evolution both by monoderm and diderm bacteria and is expected to increase the functional versatility of T4P.

**Fig. 9.**
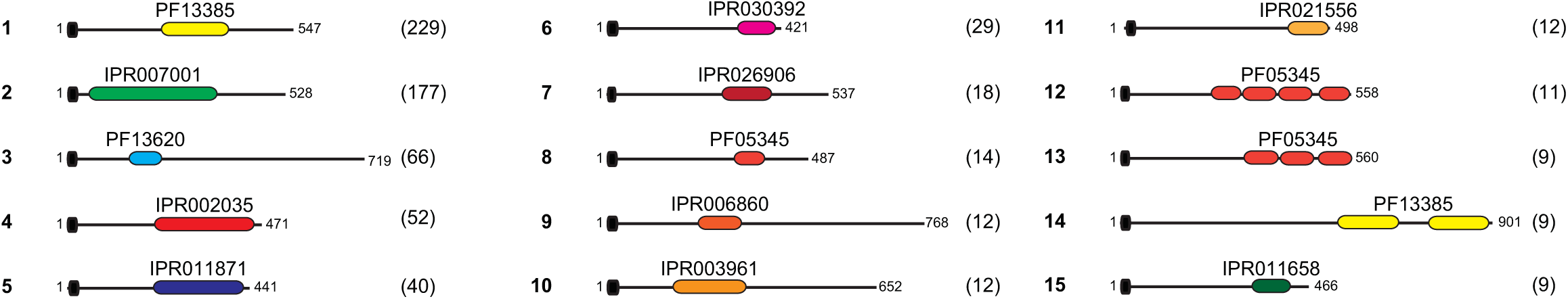
Global distribution of modular pilins. The fifteen most widespread modular pilin architectures in the InterPro database are presented. The numbers in parenthesis represent the number of proteins displaying that architecture. The representative proteins depicted, drawn to scale, are from the following species. 1, *Candidatus* Magasanikbacteria (UniProtKB/TrEMBL protein A0A0G0IU57). 2, *Photorhabdus luminescens* (A0A022PI42). 3, *Desulfuribacillus stibiiarsenatis* (A0A1E5L295). 4, *Candidatus* Falkowbacteria (A0A1J4TDE2). 5, *Candidatus* Wolfebacteria (A0A0G1WFE5). 6, Clostridiales bacterium (A0A101H8M7). 7, *Corynebacterium glutamicum* (A0A1Q6BQB1). 8, *Thermosulfidibacter takaii* (A0A0S3QUH2). 9, *Candidatus* Gracilibacteria (A0A1J5F7A7). 10, *Candidatus* Saccharibacteria (A0A1Q3NLQ9). 11, Planctomycetes bacterium (A0A1G2ZHU9). 12, *Actinoplanes awajinensis* subsp. *mycoplanecinus* (A0A101J7V4). 13, *Actinoplanes derwentensis* (A0A1H2D7E9). 14, Desulfobacterales bacterium (A0A1V1WSE4). 15, Parcubacteria group bacterium (A0A2D6FLV5).

## DISCUSSION

T4F are an important research topic because of their virtual ubiquity in prokaryotes and their ability to mediate several key biological processes^1^. Furthermore, the molecular mechanisms of most T4F-mediated functions and the exact role of minor pilins remain incompletely understood. Therefore, in this report we focused on T4aP – the prototypical T4F^1^ – in the recently established monoderm model *S. sanguinis*^15^, and performed a structure/function analysis of the unusual minor pilin PilB, which we predicted might play a role in T4P-mediated adhesion. This led to several notable findings discussed below, and confirmed predictions that the study of T4P in monoderms has the potential to shine new light on these filaments^14,15^.

The first important finding in this study is that PilB defines a new class of widespread and extremely diverse type IV pilins – the modular pilins – in which an N-terminal pilin module is fused via a short linker to distinct modules that mediate clearly defined functions. Modular pilins are likely to be tip-exposed in the filaments because of their peculiar architecture. While previous 3D structures of a few large minor pilins suggest that they are modular pilins, their second modules have no clear function and do not correspond to protein domains identifiable by available bioinformatic tools. In CofB from enterotoxigenic *E. coli* (ETEC) T4bP, there are two additional structural domains linked to the C-terminus of the pilin module by a flexible linker, a β-repeat domain followed by a β-sandwich domain^30^. CofB, which forms a trimer predicted to be exposed at the tip of ETEC T4bP^31^, appears to be an adapter for a secreted protein CofJ^32^, having a direct role in adhesion. In ComZ from *Thermus thermophilus* T4aP, the additional structural domain is a large β-solenoid inserted not at the end of the pilin module but into the β-sheet^33^. The role of ComZ is not known but it might be involved in binding extracellular DNA to promote its uptake during transformation^33^. This modular architecture is not restricted to T4P as it is also observed for a minor pilin from another T4F, GspK from type II secretion systems (T2SS)^34^. GspK, in which the additional structural domain is an α-domain of unknown function inserted into the β-sheet of the pilin module, has also been proposed to be at the tip of T2SS pseudopili, together with two other non-modular minor pilins (GspI and GspJ) with which it interacts to form a heterotrimer^34^. These examples suggest that we have here probably underestimated the global distribution of modular pilins, which are likely to be much more widespread because in many of them the additional modules are not yet defined by protein signatures in the databases. However, what is clear from our global analysis is that the functions associated with these modular pilins are potentially extremely diverse. Although a “common theme” appears to promote the interaction of T4F with a variety of ligands – including proteins (via vWA in PilB, and the β-repeat/β-sandwich module in CofB), carbohydrates (via a variety of lectin domains including the concanavalin A-like lectin/glucanase domain in PilC), or DNA (the putative role of the β-solenoid module in ComZ) – other functions are possible. This is suggested by the modular architectures IPR012902-IPR030392 or IPR012902-IPR011493 in which the second module is a predicted peptidase belonging to S74 and M26 families, respectively.

The functional characterisation of the vWA module in PilB including its MIDAS, showing that it is a *bona fide* adhesin, is another significant achievement of this study. First, the vWA domain, which is ubiquitous in the three domains of life and has been extensively studied in eukaryotes^18,21^, has been much less studied in bacteria. Second, T4P-mediated adhesion remains among the most poorly understood T4P functions^1^. Our functional analysis of the vWA module in PilB significantly extends what was known for prokaryotic vWA-containing proteins and highlights important similarities and differences with extensively studied prokaryotic vWA-containing proteins. Our 3D structure shows that the vWA module in PilB exhibits striking similarity to the vWA domain in eukaryotic vWA-containing proteins^20,24^, with a canonical MIDAS coordinating a metal. The main difference is that the MIDAS in PilB is flanked by two protruding arms, similarly to what has been described for RrgA from *S. pneumoniae*^26^. Interestingly, RrgA is a subunit with intrinsic adhesive properties^35^of sortase-assembled pili in monoderms^36^, which are unrelated to T4P. The parallel between RrgA and PilB denotes a case of convergent evolution in which two unrelated types of pili have independently evolved a similar strategy to mediate adhesion. Testing metal binding by the MIDAS in PilB, which was previously done only for eukaryotic vWA-containing proteins^37^, highlight important similarities. MIDAS show no significant binding to Ca^2+^, and a slight preference for Mn^2+^ over Mg^2+^, although the difference in affinity is much smaller than in eukaryotic proteins^37^. Metal binding can be abolished by altering the MIDAS motif, which has no impact on PilB structure^37,38^. Abolishing metal binding has no detectable effect on piliation, which is analogous to what has been reported for vWA-containing adhesins of sortase-assembled pili^39,40^, but it impairs T4P-mediated twitching motility. It is unclear at this stage whether the lack of motility of the *pilB*_*D319A*_ mutant is due to a reduced T4P adhesion to the agar, which would be consistent with PilB role in adhesion, or to impaired filament retraction, which powers movement^11^. We also provide evidence that the vWA module of PilB binds several human protein ligands, which it shares with eukaryotic vWA-containing proteins such as integrins and/or vWF^19,21^. However, unlike in these proteins where binding is often impaired when the MIDAS is inactivated^20^, a notable difference is that binding to fibronectin and fibrinogen is unaffected in a PilB_D319A_ mutant. This either suggests that the MIDAS is not implicated in binding these specific ligands, which has been described for vWF binding to collagen^25^, or that our *in vitro* binding assays are not sensitive enough to detect subtle but significant differences in binding.

The finding that PilB plays a key role in adhesion of *S. sanguinis* to host cells and structures, via its vWA module, has implications for the pathogenesis of this species in particular, and for our understanding of T4P-mediated adhesion in general. Our findings are consistent with the possibility that PilB-mediated adhesion to host proteins might play a role in IE^41^, a life-threatening infection often caused by *S. sanguinis*. Indeed, during IE, bacteria that have gained access to the bloodstream adhere to pre-existing sites of valvular damage where sub-endothelial ECM proteins are exposed, and a blood clot is present, containing large amounts of platelets, fibrinogen/fibrin, and fibronectin^42^. Our finding that PilB adheres directly to two of these proteins, but additional ligands cannot be excluded, suggests that PilB might be important at this early stage in IE, which could be tested in future studies. Our findings, which arguably make PilB the best characterised T4P adhesin alongside PilC/PilY1 found in diderm T4aP^43–47^, have general implications for our understanding of T4P-mediated adhesion. The vWA module in PilB, which is most likely exposed at the pilus tip, is ideally placed to maximise bacterial adhesion to host protein receptors. T4P spring-like properties (gonococcal T4P can be stretched 3 times their length, which is a reversible process^48^) are expected to help bacteria that are bound via a tip-located adhesin, such as PilB, to withstand adverse forces in their particular environment, *e*.*g*., blood flow in a heart valve. This is likely to apply to other modular pilins as well, which harbour different modules predicted to function in adhesion. The parallel with the best characterised T4P adhesin PilC/PilY1 is obvious. This protein, which is not a pilin, is an adhesin that has been proposed to be presented at the T4P tip^44^ via its interaction with a tip-located complex of four widely conserved minor pilins^49^. All PilC/PilY1 have in common a C-terminal IPR008707 β-propeller domain while their N-termini are different^50^. Since this is analogous to the situation with modular pilins, we wondered whether it could be an indication of a modular design for PilC/PilY1. This indeed seems to be the case since a search of the InterPro database^29^ for all the proteins with an IPR008707 domain shows that 68 different PilC/PilY1 modular architectures are detected (Supplementary data 2). Strikingly, many of the N-terminal modules in PilC/PilY1 are shared with modular pilins. These observations suggest that the same tinkering strategy has been used both by pilins and PilC/PilY1 to increase the functional versatility of T4P. In both instances, a “carrier” module for presentation at the tip of the filaments (either a pilin, or an IPR008707 domain that interacts with a tip-located complex of minor pilins) has been fused to variety of “effector” modules, directly involved in a variety of functions.

In conclusion, by performing a detailed structure/function of the minor pilin PilB from *S. sanguinis*, we have shed light on several aspects of T4P biology. Our findings are not only of relevance for *S. sanguinis*, most notably for colonisation of its human host, they have general implications for T4P/T4F by uncovering a prevalent strategy used by these widespread filamentous nanomachines to promote their well-known exceptional functional versatility^1^. The resulting conceptual framework paves the way for further investigations, which will indubitably improve our understanding of theses fascinating filaments.

## MATERIALS AND METHODS

### Strains and growth conditions

Strains and plasmids used in this study are listed in Table S1. For cloning, we used *E. coli* DH5α. For protein purification we used *E. coli* BL21(DE3) or *E. coli* BL21 B834(DE3). *E. coli* was grown in liquid or solid Lysogenic Broth (LB) (Difco) containing, when required, 100 µg/ml spectinomycin or 50 µg/ml kanamycin (both from Sigma). For purification of protein labelled with seleno-methionine (SeMet), bacteria were grown in chemically defined medium (CDM) supplemented with 20 mg/ml SeMet (Sigma). Chemically competent *E. coli* cells were prepared as described^51^. DNA manipulations were done using standard molecular biology techniques^52^. All PCR were done using high-fidelity DNA polymerases (Agilent). Primers used in this study are listed in Table S2. The pET-28b (Novagen) derivative, pET28-*pilB*_*36-461*_ for expressing 6His-PilB_36-461_ was described previously^17^. In this plasmid, the portion of a synthetic *pilB* gene codon-optimised for expression in *E. coli*, encoding the soluble portion of PilB, was fused to a non-cleavable N-terminal 6His tag. Similarly, we constructed pET28-*pilB*_*192-461*_ for expressing PilB_vWA_. To construct pET28-*pilB*_*D319A*_ for expressing 6His-PilB_D319A_, we introduced a missense mutation in pET28-*pilB*_*36-461*_ using QuickChange site-directed mutagenesis (Agilent).

The WT *S. sanguinis* 2908 strain and deletion mutants (*ΔpilD, ΔpilB*) were described previously^16^. *S. sanguinis* was grown on plates containing Todd Hewitt (TH) broth (Difco) and 1% agar (Difco), incubated at 37°C in anaerobic jars (Oxoid) under anaerobic conditions generated using Anaerogen sachets (Oxoid). Liquid cultures were grown statically under aerobic conditions in THT, *i*.*e*., TH broth containing 0.05% tween 80 (Merck) to limit bacterial clumping. When required, 500 μg/ml kanamycin was used for selection and 15 mM *p*-Cl-Phe (Sigma) for counterselection^53^. *S. sanguinis* genomic DNA was prepared from overnight (O/N) liquid cultures using the kit XIT Genomic DNA from Gram-Positive Bacteria (G-Biosciences). Strain 2908, which is naturally competent, was transformed as described^16,53^. The unmarked *S. sanguinis pilB*_*D319A*_ mutant was constructed using a previously described two-step, cloning-independent, gene editing strategy^53^. In brief, in the first step, we replaced the gene in the WT by a promoterless *pheS*aphA-3* double cassette, which confers sensitivity to *p*-Cl-Phe and resistance to kanamycin. To do this, we fused by splicing PCR the upstream and downstream regions flanking *pilB* to *pheS*aphA-3*, directly transformed the PCR product into the WT, and selected allelic exchange mutants on kanamycin plates. Allelic exchange was confirmed by PCR. In the second step, we replaced the *pheS*aphA-3* double cassette in this primary mutant by allelic exchange, with an unmarked *pilB*_*D319A*_ mutation. To do this, we first constructed the missense mutation by site-directed mutagenesis, using as a template a pCR8/GW/TOPO (Invitrogen) derivative in which the WT gene was cloned. Then, the PCR product was directly transformed into the primary mutant, with plating on *p*-Cl-Phe-containing plates. Markerless allelic exchange mutants, which are the only one sensitive to kanamycin, were identified by re-streaking colonies on TH plates with and without antibiotic.

### Protein purification

To purify native PilB, PilB_D319A_ and PilB_vWA_ proteins, the corresponding pET-28b derivatives were transformed in *E. coli* BL21(DE3). Transformants were grown O/N at 37°C in liquid LB with kanamycin. The next day, this culture was diluted (1/500) in 1 l of the same medium and grown at 37°C to an OD_600_ of 0.4-0.6. The temperature was then set to 16°C, the culture allowed to cool for 30 min, before protein expression was induced O/N by adding 0.5 mM IPTG (Merck). The next day, cells were harvested by centrifugation at 8,000 *g* for 20 min and subjected to one −80°C freeze/thaw cycle in binding buffer (50 mM HEPES pH 7.4, 200 mM NaCl, 15 mM imidazole), to which we added SIGMAFAST EDTA-free protease inhibitor cocktail (Sigma). Cells were disrupted by repeated cycles of sonication, *i*.*e*., pulses of 5 sec on and 5 sec off during 3-5 min, until the cell suspension was visibly less viscous. The cell lysate was then centrifuged for 30 min at 17,000 *g* to remove cell debris. The clarified lysate was first affinity-purified on an ÄKTA Purifier using His-Trap HP columns (GE Healthcare) and eluted with elution buffer (50 mM HEPES pH 7.4, 200 mM NaCl, 300 mM imidazole). Affinity-purified proteins were further purified, and simultaneously buffer-exchanged into (50 mM HEPES pH 7.4, 200 mM NaCl), by gel-filtration chromatography on an ÄKTA Purifier using a Superdex 75 10/300 GL column (GE Healthcare). Protein concentration was quantified spectrophotometrically using a NanoDrop.

To purify SeMet-labelled PilB for phasing, the corresponding pET-28b derivative was transformed in *E. coli* BL21 B834(DE3). Transformants were grown at 37°C in liquid LB with kanamycin, until OD_600_ reached 0.6-0.7. Next, the cells were pelleted at 8,000 *g* for 5 min, and washed twice with 2 ml of CDM, which contains no Met. The pellet was then washed with 2 ml of CDM supplemented with 20 mg/ml L-Met (Sigma) and used to inoculate, at 1/200 dilution, 20 ml of CDM supplemented with 20 mg/ml Met. This culture was grown at 37°C for 16-18 h. Cells were pelleted and washed three times with CDM. Then, the pellet was re-suspended in 20 ml of CDM supplemented with 20 mg/ml SeMet, which was used to inoculate 1 l of CDM supplemented with SeMet. Cells were grown at 37°C until OD_600_ reached 0.5-0.7. The temperature was then set to 16°C, the culture allowed to cool for 30 min, before protein expression was induced O/N by adding 1 mM IPTG (Merck) and 4 ml of 36 % glucose (w/v). Two and half hours later, we again added 4 ml of 36 % glucose to the culture. The next day, SeMet-labelled PilB was purified as above.

### Crystallisation and structure determination

Purified proteins in 50 mM HEPES pH 7.4, 200 mM NaCl were concentrated to 50 mg/ml and tested for crystallisation using sitting-drop vapor diffusion, with 100 nl drops of protein solution and mother liquor. We tested a range of commercially available kits (Molecular Dimensions, Hampton Research and Rigaku Reagents), which yielded a number of hits, mainly in high salt conditions. Crystallisation conditions were optimised to yield larger and better diffracting crystals. The PilB crystals used for the high-resolution structure determination were obtained when the purified protein was mixed 1:1 with crystallisation liquor (0.1 M Bis-tris propane pH 7, 3 M NaCl). The PilB_D319A_ crystals were obtained with crystallisation liquor (0.1 M Bis-tris pH 6.5, 3 M NaCl). Crystals were cryoprotected with 30% glycerol in crystallisation liquor, and flash-frozen in liquid nitrogen. All data was collected and processed using the Diamond Light Source beamline i03, and integrated in *P*6_1_ using the 3dii pipeline in *xia*2^54^. Initial molecular replacement was performed with Phaser^55^ on the 2.26 Å resolution PilB dataset using a low-resolution partial model produced from the SeMet data using autoSHARP^56^. Manual building in Coot^57^ was performed on the high-resolution dataset, and the full model was then used for molecular replacement in the low-resolution datasets. All structures were produced using Coot and phenix.refine^58^ and validated using MolProbity^59^.

### Assaying metal-binding by purified PilB

The metal binding specificity of PilB was tested using ThermoFluor, a fluorescent-based method measuring changes in thermal denaturation temperature^27^. Assays were done in a 96-well plate (Applied Biosystems) format. In the wells, we added to a final volume of 40 µl (i) 0-1 mM range of concentrations of MgCl_2_, MnCl_2_ and CaCl_2_, (ii) 20 µM purified PilB or PilB_D319A_, and (iii) 1/5,000 dilution of SYBR Orange (Thermo Fisher Scientific). Plates were then analysed using a temperature gradient, from 25 to 99 °C, on a StepOnePlus real-time qPCR machine (Applied Biosystems). The data were exported MATLAB and analysed in GraphPad. Analyses were performed with Prism (GraphPad Software). Kd were calculated using non-linear regression fits, applying saturation binding equation (One site - Total and non-specific binding) using Ca^2+^ as non-specific binding control.

### Assaying protein ligand-binding by purified PilB

Binding of PilB, PilB_vWA_, PilB_D319A_ to a variety of eukaryotic proteins was tested by ELISA as follows. Putative ligand proteins (elastin from human skin, fibrinogen from human plasma, laminin from human placenta, fibronectin from human plasma) (all from Sigma) were resuspended in carbonate-bicarbonate buffer (Sigma) at 5 µg/ml. Fifty µl was dispatched into the wells of MaxiSorp plates, and adsorbed O/N at 4°C. Wells were washed three times with PBS (Gibco) and blocked during 1 h with 3 % BSA (Probumin) or 1% gelatin (Sigma) in PBS. After washing with PBST (PBS containing 0.05 % tween 20), serial twofold dilutions of PilB (from 40 to 0.625 µg/ml) were added to the wells and incubated for 2 h at 37°C. After five washes with PBST, we added 50 µl anti-6His RTM antibody (Abcam) at 1/500 dilution in PBS, and incubated for 1 h at RT. After five washes with PBST, we added 50 µl Amersham ECL anti-rabbit IgG HRP-linked whole antibody (GE Healthcare) at 1/500 dilution in PBS, and incubated for 1 h at RT. After five washes with PBST, we added 100 µl/well of TMB solution (Thermo Scientific) and incubated the plates during 20 min at RT in the dark. Finally, we stopped the reaction by adding 100 µl/well of 0.18 M sulfuric acid, before reading the plates at 450 nm using a plate reader. Analyses were performed with Prism (GraphPad Software). Kd were calculated using non-linear regression fits, applying saturation binding equation (One site - Total and non-specific binding) using BSA or gelatin as non-specific binding control.

### Assaying piliation of *S. sanguinis*

*S. sanguinis* T4P were purified as described^16,17^. In brief, bacteria grown in THT until the OD_600_ reached 1-1.5, at which point OD were normalised, a were pelleted and re-suspended in pilus buffer (20 mM Tris, pH 7.5, 50 mM NaCl). This suspension was vortexed for 2 min at full speed to shear T4P. After removing bacterial cells by two centrifugation steps and filtration through a 0.22 µm pore size syringe filter (Millipore), pili were pelleted by ultracentrifugation. Pili were resuspended in pilus buffer, separated by SDS-PAGE, before gels were stained with Bio-Safe Coomassie (Bio-Rad).

### Assaying twitching motility of *S. sanguinis*

Twitching motility was assessed on agar plates as described^16^. In brief, bacteria grown O/N were streaked as straight lines on freshly poured TH plates containing 1% Eiken agar (Eiken Chemicals). Plates were grown for several days at 37°C in anaerobic condition under high humidity, which is necessary for twitching. Plates were then photographed using an Epson Perfection V700 photo scanner.

### Assaying adhesion of *S. sanguinis* to eukaryotic cells

We tested adhesion of *S. sanguinis* to CHO cells (Public Health England) as follows. Cells were replicated in flasks in DMEM medium (Gibco) containing 1 × MEM non-essential aa mix (Gibco) and 5 % fetal bovine serum (Gibco) and seeded at 100,000 cells/cm^2^ in 24-well plates, which were incubated O/N at 37°C in the presence of 5% CO_2_. The next day, cell monolayers were gently rinsed with PBS, and infected at an MOI of 10 with bacteria grown in TH. In brief, bacteria were grown for a few hours to OD_600_ 0.5 units, adjusted at the same OD, pelleted by centrifugation at 1,100 *g* during 10 min, and resuspended in PBS. Bacteria in the inoculum were quantified by performing CFU counts on TH plates. After 1h of infection at 37°C, cell monolayers were gently rinsed four times with PBS, before cells with adherent bacteria were scraped in distilled water. Adherent bacteria were then quantified by performing CFU counts. Statistical analyses were performed with Prism. Comparisons were done by one-way ANOVA, followed by Dunnett’s multiple comparison tests. An adjusted *P* value < 0.05 was considered significant (* *P*<0.05, ** *P*<0.01, *** *P*<0.001, **** *P*<0.0001).

### Bioinformatics

Protein sequences were routinely analysed using DNA Strider^60^. Prediction of protein domains, their global distribution and associated architectures was done by using InterProScan^29^ to interrogate the InterPro database. This database was also used to download all the protein entries discussed in this paper. Molecular visualisation of protein 3D structures was done using PyMOL (Schrödinger). The PDBsum Generate^61^ server was used to provide at-a-glance overviews – secondary structure, topology diagram, protein motifs, schematic diagram of metal-protein interactions – of the 3D structures determined during this work. The DALI^62^ server was used for comparing protein structures in 3D. Protein 3D structures were downloaded from the RCSB PDB server. The 3d-SS^63^ server was used to superpose 3D protein structures.

The cryo-EM structure of *N. meningitidis* T4P (PDB 5KUA)^9^ was used to model, using SWISS-MODEL^64^, the N-terminal helices of PilE1, PilE2 and PilB within the filaments. Coot and PyMOL were then used to place the full-length structures within the T4P model.

## DATA AVAILABILITY

The 3D structures determined during this study have been deposited in the PDB, under accession codes 7B7P for PilB, and 7BA2 for PilB_D319A_. All the datasets generated during this study are included in this paper and its Supplementary information. Source data are provided. The InterPro database (http://www.ebi.ac.uk/interpro) was used to identify modular pilin architectures. The DALI server (http://ekhidna2.biocenter.helsinki.fi/dali) was used for comparing protein 3D structures. The PDBsum Generate server (https://www.ebi.ac.uk/thornton-srv/databases/pdbsum/Generate.html) was used to generate analytical overviews of the 3D structures we have determined. The 3d-SS server (http://cluster.physics.iisc.ernet.in/3dss) was used for 3D superposition of protein structures. The RCSB PDB server (https://www.rcsb.org) was used to download x3D structures of proteins.

## ACKNOWLEDGEMENTS

This work was supported by funding from the MRC (MR/P022197/1) to V. P. We acknowledge the use of the crystallisation facility at Imperial College London, which was supported by BBSRC (BB/D524840/1) and the Wellcome Trust (202926/Z/16/Z). We thank Angelika Gründling (Imperial College London), Sophie Helaine (Harvard Medical School), Christoph Tang (University of Oxford), and Romé Voulhoux (Laboratoire de Chimie Bactérienne) for critical reading of the manuscript.

## AUTHOR CONTRIBUTIONS

V. P. was responsible for conception and supervision of the work, and the computational studies. C. R., D. S., J. L. B., and I. G. performed the experimental studies. All authors contributed to writing of the manuscript

## REFERENCES

1. Berry, J.L. & Pelicic, V. Exceptionally widespread nano-machines composed of type IV pilins: the prokaryotic Swiss Army knives. FEMS Microbiology Reviews 39, 134–154 (2015).

2. Denise, R., Abby, S.S. & Rocha, E.P.C. Diversification of the type IV filament superfamily into machines for adhesion, protein secretion, DNA uptake, and motility. PLoS Biology 17, e3000390 (2019).

3. Pelicic, V. Type IV pili: e pluribus unum? Molecular Microbiology 68, 827–837 (2008).

4. Giltner, C.L., Nguyen, Y. & Burrows, L.L. Type IV pilin proteins: versatile molecular modules. Microbiology and Molecular Biology Reviews 76, 740–72 (2012).

5. LaPointe, C.F. & Taylor, R.K. The type 4 prepilin peptidases comprise a novel family of aspartic acid proteases. Journal of Biological Chemistry 275, 1502–10 (2000).

6. Francetic, O., Buddelmeijer, N., Lewenza, S., Kumamoto, C.A. & Pugsley, A.P. Signal recognition particle-dependent inner membrane targeting of the PulG pseudopilin component of a type II secretion system. Journal of Bacteriology 189, 1783–93 (2007).

7. Arts, J., van Boxtel, R., Filloux, A., Tommassen, J. & Koster, M. Export of the pseudopilin XcpT of the Pseudomonas aeruginosa type II secretion system via the signal recognition particle-Sec pathway. Journal of Bacteriology 189, 2069–76 (2007).

8. Chang, Y.W. et al. Architecture of the type IVa pilus machine. Science 351, aad2001 (2016).

9. Kolappan, S. et al. Structure of the Neisseria meningitidis type IV pilus. Nature Communications 7, 13015 (2016).

10. Wang, F. et al. Cryoelectron microscopy reconstructions of the Pseudomonas aeruginosa and Neisseria gonorrhoeae type IV pili at sub-nanometer resolution. Structure 25, 1423–1435 (2017).

11. Merz, A.J., So, M. & Sheetz, M.P. Pilus retraction powers bacterial twitching motility. Nature 407, 98–102 (2000).

12. Maier, B. et al. Single pilus motor forces exceed 100 pN. Proceedings of the National Academy of Sciences of the United States of America 99, 16012–7 (2002).

13. Biais, N., Ladoux, B., Higashi, D., So, M. & Sheetz, M. Cooperative retraction of bundled type IV pili enables nanonewton force generation. PLoS Biology 6, e87 (2008).

14. Melville, S. & Craig, L. Type IV pili in Gram-positive bacteria. Microbiology and Molecular Biology Reviews 77, 323–41 (2013).

15. Pelicic, V. Monoderm bacteria: the new frontier for type IV pilus biology. Molecular Microbiology 112, 1674–1683 (2019).

16. Gurung, I. et al. Functional analysis of an unusual type IV pilus in the Gram-positive Streptococcus sanguinis. Molecular Microbiology 99, 380–392 (2016).

17. Berry, J.L. et al. Global biochemical and structural analysis of the type IV pilus from the Gram-positive bacterium Streptococcus sanguinis. Journal of Biological Chemistry 294, 6796–6808 (2019).

18. Whittaker, C.A. & Hynes, R.O. Distribution and evolution of von Willebrand/integrin A domains: widely dispersed domains with roles in cell adhesion and elsewhere. Molecular Biology of the Cell 13, 3369–87 (2002).

19. Sadler, J.E. Biochemistry and genetics of von Willebrand factor. Annual Review of Biochemistry 67, 395–424 (1998).

20. Lee, J.O., Rieu, P., Arnaout, M.A. & Liddington, R. Crystal structure of the A domain from the alpha subunit of integrin CR3 (CD11b/CD18). Cell 80, 631–8 (1995).

21. Dickeson, S.K. & Santoro, S.A. Ligand recognition by the I domain-containing integrins. Cellular and Molecular Life Sciences 54, 556–66 (1998).

22. Craig, L. et al. Type IV pilin structure and assembly. X-ray and EM analyses of Vibrio cholerae toxin-coregulated pilus and Pseudomonas aeruginosa PAK pilin. Molecular Cell 11, 1139–50 (2003).

23. Hutchinson, E.G. & Thornton, J.M. HERA–a program to draw schematic diagrams of protein secondary structures. Proteins 8, 203–12 (1990).

24. Qu, A. & Leahy, D.J. Crystal structure of the I-domain from the CD11a/CD18 (LFA-1, αLβ2) integrin. Proceedings of the National Academy of Sciences of the United States of America 92, 10277–81 (1995).

25. Romijn, R.A. et al. Identification of the collagen-binding site of the von Willebrand factor A3-domain. Journal of Biological Chemistry 276, 9985–91 (2001).

26. Izoré, T. et al. Structural basis of host cell recognition by the pilus adhesin from Streptococcus pneumoniae. Structure 18, 106–15 (2010).

27. Niesen, F.H., Berglund, H. & Vedadi, M. The use of differential scanning fluorimetry to detect ligand interactions that promote protein stability. Nature Protocols 2, 2212–21 (2007).

28. Kreth, J., Giacaman, R.A., Raghavan, R. & Merritt, J. The road less traveled - defining molecular commensalism with Streptococcus sanguinis. Molecular Oral Microbiology 32, 181–196 (2017).

29. Jones, P. et al. InterProScan 5: genome-scale protein function classification. Bioinformatics 30, 1236–40 (2014).

30. Kolappan, S., Ng, D., Yang, G., Harn, T. & Craig, L. Crystal structure of the minor pilin CofB, the initiator of CFA/III pilus assembly in enterotoxigenic Escherichia coli. Journal of Biological Chemistry 290, 25805–18 (2015).

31. Kawahara, K. et al. Homo-trimeric structure of the type IVb minor pilin CofB suggests mechanism of CFA/III pilus assembly in human enterotoxigenic Escherichia coli. Journal of Molecular Biology 428, 1209–1226 (2016).

32. Oki, H. et al. Interplay of a secreted protein with type IVb pilus for efficient enterotoxigenic Escherichia coli colonization. Proceedings of the National Academy of Sciences of the United States of America (2018).

33. Salleh, M.Z. et al. Structure and properties of a natural competence-associated pilin suggest a unique pilus tip-associated DNA receptor. mBio 10, e00614–19 (2019).

34. Korotkov, K.V. & Hol, W.G. Structure of the GspK-GspI-GspJ complex from the enterotoxigenic Escherichia coli type 2 secretion system. Nature Structural and Molecular Biology 15, 462–8 (2008).

35. Hilleringmann, M. et al. Pneumococcal pili are composed of protofilaments exposing adhesive clusters of RrgA. PLoS Pathogens 4, e1000026 (2008).

36. Telford, J.L., Barocchi, M.A., Margarit, I., Rappuoli, R. & Grandi, G. Pili in Gram-positive pathogens. Nature Reviews in Microbiology 4, 509–19 (2006).

37. Baldwin, E.T. et al. Cation binding to the integrin CD11b I domain and activation model assessment. Structure 6, 923–35 (1998).

38. Qu, A. & Leahy, D.J. The role of the divalent cation in the structure of the I domain from the CD11a/CD18 integrin. Structure 4, 931–42 (1996).

39. Konto-Ghiorghi, Y. et al. Dual role for pilus in adherence to epithelial cells and biofilm formation in Streptococcus agalactiae. PLoS Pathogens 5, e1000422 (2009).

40. Nielsen, H.V. et al. The metal ion-dependent adhesion site motif of the Enterococcus faecalis EbpA pilin mediates pilus function in catheter-associated urinary tract infection. MBio 3, e00177–12 (2012).

41. Que, Y.A. & Moreillon, P. Infective endocarditis. Nature Reviews Cardiology 8, 322–36 (2011).

42. Werdan, K. et al. Mechanisms of infective endocarditis: pathogen-host interaction and risk states. Nature Reviews Cardiology 11, 35–50 (2014).

43. Nassif, X. et al. Roles of pilin and PilC in adhesion of Neisseria meningitidis to human epithelial and endothelial cells. Proceedings of the National Academy of Sciences of the United States of America 91, 3769–3773 (1994).

44. Rudel, T., Scheuerpflug, I. & Meyer, T.F. Neisseria PilC protein identified as a type-4 pilus-tip located adhesin. Nature 373, 357–359 (1995).

45. Heiniger, R.W., Winther-Larsen, H.C., Pickles, R.J., Koomey, M. & Wolfgang, M.C. Infection of human mucosal tissue by Pseudomonas aeruginosa requires sequential and mutually dependent virulence factors and a novel pilus-associated adhesin. Cellular Microbiology 12, 1158–73 (2010).

46. Johnson, M.D. et al. Pseudomonas aeruginosa PilY1 binds integrin in an RGD- and calcium-dependent manner. PLoS One 6, e29629 (2011).

47. Porsch, E.A. et al. Calcium binding properties of the Kingella kingae PilC1 and PilC2 proteins have differential effects on type IV pilus-mediated adherence and twitching motility. Journal of Bacteriology 195, 886–95 (2013).

48. Biais, N., Higashi, D.L., Brujic, J., So, M. & Sheetz, M.P. Force-dependent polymorphism in type IV pili reveals hidden epitopes. Proceedings of the National Academy of Sciences of the United States of America 107, 11358–63 (2010).

49. Treuner-Lange, A. et al. PilY1 and minor pilins form a complex priming the type IVa pilus in Myxococcus xanthus. Nature Communications 11, 5054 (2020).

50. Orans, J. et al. Crystal structure analysis reveals Pseudomonas PilY1 as an essential calcium-dependent regulator of bacterial surface motility. Proceedings of the National Academy of Sciences of the United States of America 107, 1065–70 (2010).

51. Inoue, H., Nojima, H. & Okayama, H. High efficiency transformation of Escherichia coli with plasmids. Gene 96, 23–8 (1990).

52. Sambrook, J. & Russell, D.W. Molecular cloning. A laboratory manual, (Cold Spring Harbor Laboratory Press, Cold Spring Harbor, New York, 2001).

53. Gurung, I., Berry, J.L., Hall, A.M.J. & Pelicic, V. Cloning-independent markerless gene editing in Streptococcus sanguinis: novel insights in type IV pilus biology. Nucleic Acids Research 45, e40 (2017).

54. Winter, G., Lobley, C.M. & Prince, S.M. Decision making in xia2. Acta Crystallographica Section D Biological Crystallography 69, 1260–73 (2013).

55. McCoy, A.J. et al. Phaser crystallographic software. Journal of Applied Crystallography 40, 658–674 (2007).

56. Vonrhein, C., Blanc, E., Roversi, P. & Bricogne, G. Automated structure solution with autoSHARP. Methods in Molecular Biology 364, 215–30 (2007).

57. Emsley, P., Lohkamp, B., Scott, W.G. & Cowtan, K. Features and development of Coot. Acta Crystallographica Section D Biological Crystallography 66, 486–501 (2010).

58. Afonine, P.V. et al. Towards automated crystallographic structure refinement with phenix.refine. Acta Crystallographica Section D Biological Crystallography 68, 352–67 (2012).

59. Chen, V.B. et al. MolProbity: all-atom structure validation for macromolecular crystallography. Acta Crystallographica Section D Biological Crystallography 66, 12–21 (2010).

60. Marck, C. ‘DNA Strider’: a ‘C’ program for the fast analysis of DNA and protein sequences on the Apple Macintosh family of computers. Nucleic Acids Research 16, 1829–36 (1988).

61. Laskowski, R.A., Jablonska, J., Pravda, L., Varekova, R.S. & Thornton, J.M. PDBsum: Structural summaries of PDB entries. Protein Science 27, 129–134 (2018).

62. Holm, L. & Laakso, L.M. Dali server update. Nucleic Acids Research 44, W351–5 (2016).

63. Sumathi, K., Ananthalakshmi, P., Roshan, M.N. & Sekar, K. 3dSS: 3D structural superposition. Nucleic Acids Research 34, W128–32 (2006).

64. Waterhouse, A. et al. SWISS-MODEL: homology modelling of protein structures and complexes. Nucleic Acids Research 46, W296–W303 (2018).

