## Supplementary information for "PilB from *Streptococcus sanguinis* is a bimodular type IV pilin with a direct role in adhesion"

**Table S1. Strains and plasmids used in this study.**

| Name | Details | Source |
|---|---|---|
| ***E. coli* strains** | | |
| DH5α | used for cloning | |
| BL21(DE3) | used for protein expression/purification | |
| BL21 B834(DE3) | used for SeMet protein expression/purification | |
| ***S. sanguinis* strains** | | |
| 2908 | sequenced WT isolate | 16 |
| *ΔpilB* | *ΔpilB::aphA-3* deletion mutant | 16 |
| *ΔpilD* | *ΔpilD::aphA-3* deletion mutant | 16 |
| *ΔpilB* primary mutant | *ΔpilB::pheS*aphA-3* deletion mutant | this study |
| $\mathit{pilB}_{D319A}$ | *pilB* point mutant expressing $\mathrm{PilB}_{D319A}$ | this study |
| **Plasmids** | | |
| pCR8/GW/TOPO | TA cloning vector | Invitrogen |
| TOPO-*pheS*aphA-3* | *pheS*aphA-3* double cassette in TOPO | 53 |
| TOPO-*pilB* | *pilB* in TOPO | this study |
| TOPO-$\mathit{pilB}_{D319A}$ | $\mathit{pilB}_{D319A}$ in TOPO | this study |
| pET-28b | T7-based expression vector | Novagen |
| pMK-*pilB* | codon-optimised synthetic *pilB* in pMK | 17 |
| pET28-*pilB* | pET-28 derivative for expressing 6His-$\mathrm{PilB}_{36\text{-}461}$ | 17 |
| pET28-$\mathit{pilB}_{D319A}$ | pET-28 derivative for expressing 6His-$\mathrm{PilB}_{D319A}$ | this study |
| pET28-$\mathit{pilB}_{vWA}$ | pET-2b derivative for expressing 6His-$\mathrm{PilB}_{192\text{-}461}$ | this study |

*pilB*, codon-optimised synthetic gene. *pheS**, point mutant encoding $\mathrm{PheS}_{A316G}$.

**Table S2. Primers used in this study.**

| Name | Sequence |
|---|---|
| *pilB*$_{D319A}$ #1 | GAAATATATCGTTCTGCTGACCGcTGGTATTCCGAATGCATATCTGG |
| *pilB*$_{D319A}$ #2 | CCAGATATGCATTCGGAATACCAgCGGTCAGCAGAACGATATATTTC |
| *pilB*$_{vWA}$-pET-F | ggg**ccatgg**atcatcatcatcatcatcatCAGGGCCAGATGAATATTGC |
| *pilB*$_{vWA}$-pET-R | ccc**ggatcc**TTACGGACCGCTAACAAACC |
| *pilB-F* | TACAACTGGACCGAAGCTGG |
| *pilB-R* | TTTGGCCTATCGTTCCCACT |
| *pilB-F1* | TACAACTGGACCGAAGCTGG |
| *pilB-R1* | gttcttcaatcgttttcgtcatcaTTCCTACCTATTTATTTTTACTTCTG |
| *pilB-F2* | ttttactggatgaattgttttagAGGATTTGTGGTTTGTATCAGGG |
| *pilB-R2* | TTTGGCCTATCGTTCCCACT |
| *pheS*-F* | ATGACGAAAACGATTGAAGAAC |
| *aph-R* | CTAAAACAATTCATCCAGTAAAA |
| *pilB*$_{D319A}$ #1 | GCTTAAATATATAGTTCTATTGACAGcTGGCATACCTAATGCTTATTTAGTAG |
| *pilB*$_{D319A}$ #2 | CTACTAAATAAGCATTAGGTATGCCAgCTGTCAATAGAACTATATATTTAAGC |

*pilB*, codon-optimised synthetic gene. Overhangs are in lower case. Restriction sites are in bold. Mismatches are in red.

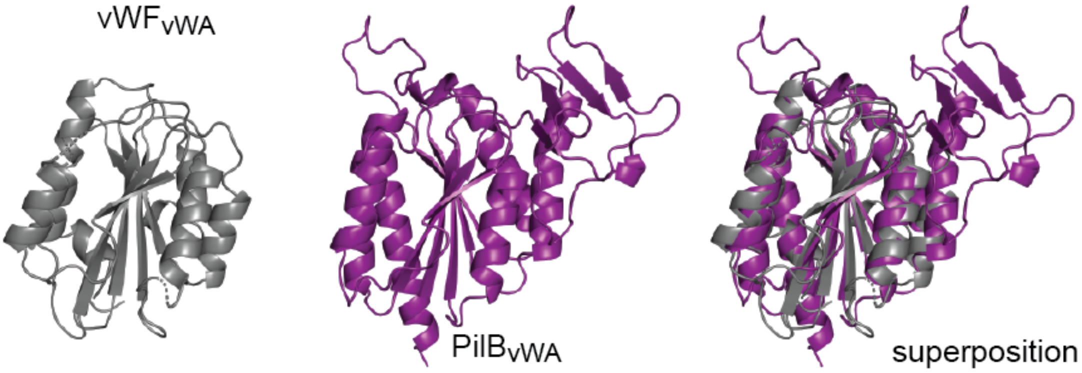


**Fig. S1. Structural similarity of vWA domains in PilB and human vWF.** Left, A3 vWA domain in vWF (from PDB 1FE8) (grey). Center, vWA module of PilB (purple). Right, superposition of the two structures. While the two sequences share only 15.6 % sequence identity, the two structures superpose with an RMSD of 1.72 Å.

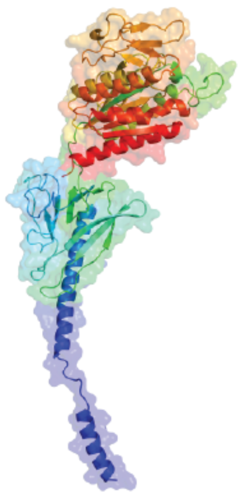

**Fig. S2. 3D model of full-length PilB with a melted α1N segment.** The cryo-EM structure of the *N. meningitidis* T4P (PDB 5KUA) has been used as a template.

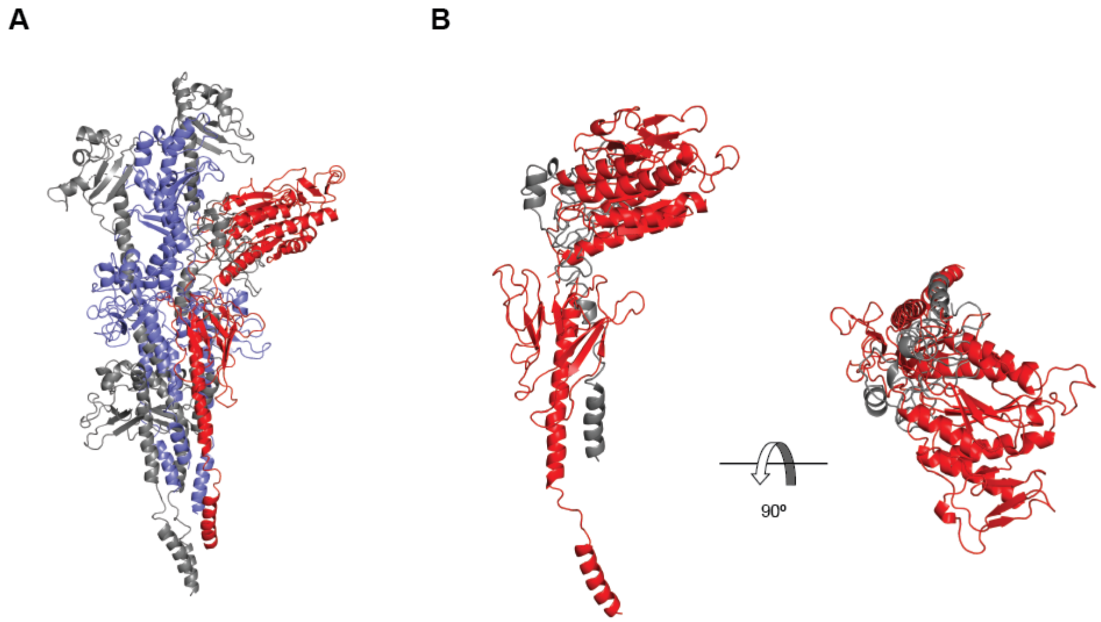


**Fig. S3. Modelling of PilB in the body of a T4P leads to important steric clashes. A**) Packing of PilB (red) into *S. sanguinis* T4P composed of PilE1 (blue) and PilE2 (grey). **B**) Close-up view of the important steric clashes between PilB and PilE subunit above in the filament.

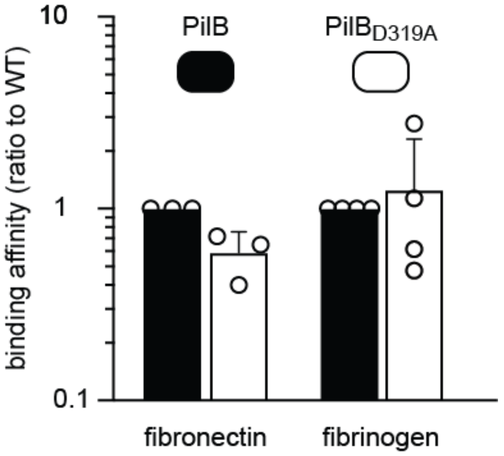


**Fig. S4. Binding of $\text{PilB}_{\text{D319A}}$ to fibrinogen and fibronectin.** Increasing concentrations of purified PilB was added to constant concentrations of immobilised ligands, and binding was quantified by ELISA. Results are represented as Kd relative to WT, which is set to 1. Results are the average ± standard deviations from 3-4 independent experiments.

**Legends to Supplementary data**

**Supplementary data 1. List of all the pilin architectures in the InterPro database.** This list was generated by searching the database (November 2020) for entries displaying an IPR012902 pilin motif. Modular pilins display an N-terminal pilin motif together with additional module(s) not specific to T4P biology. Proteins in which the pilin motif is not N-terminal are listed as unclear.

**Supplementary data 2. List of all the PilC/PilY1 architectures in the InterPro database.** This list was generated by searching the database (November 2020) for entries displaying the IPR008707 PilC/PilY1 β-propeller domain. Modular PilC/PilY1 display a C-terminal IPR008707 motif together with additional module(s) not specific to T4P biology. Proteins in which the IPR008707 motif is not C-terminal are listed as unclear.
